# Cortical neural landscape captures mouse-to-mouse variability in anticipatory vs. inattentive decision making

**DOI:** 10.1101/2025.07.11.664420

**Authors:** Chang Yin, Naoki Hiratani

## Abstract

Understanding individual variability in behavior is crucial for both basic and clinical neuroscience, yet it remains challenging to study in traditional single laboratory experiments with small sample size. Leveraging standardized behavioral and neural datasets from the International Brain Laboratory, comprising approximately 100 mice trained on a visual decision-making task, we investigated the structure and neural correlates of inter-animal behavioral variability. Using reaction time analysis and a deep learning-based embedding of individual animals, we uncovered large but low-dimensional differences in behavioral traits. Some mice consistently exhibited anticipatory responses, marked by fast reaction times, while others showed slower, more disengaged behavior. These behavioral profiles were consistent across sessions, with female mice tending to show more anticipatory behavior than males. We hypothesized that this behavioral spectrum reflects differences in the depth of underlying cortical states, reflected in the temporal dynamics of neural activity. Supporting this idea, we found that the characteristic timescale of population activity, measured during both inter-trial intervals and passive periods, correlated with an animal’s anticipatory tendency across cortical areas, especially in medial visual areas. These findings suggest that individual differences in the cortical dynamics may underlie distinct decision-making strategies.

## INTRODUCTION

Individual variability is a hallmark of social animals. Even genetically homogeneous inbred mice and rats show considerable differences in stress resistance^1–3^, task-acquisition speed^4–6^, and decision-making behavior^7,8^. Traditionally, such inter-animal variability has been viewed as a nuisance that undermines the reliability and reproducibility of experiments^9,10.^ While efforts to reduce variability have led to valuable guidelines for improving experimental rigor, variability itself can be informative rather than merely problematic^11,12.^ Natural variability offers a unique opportunity to test theories of brain function, which must ultimately account for both behavioral and neural diversity. Moreover, understanding across-individual variability in behavior and its neural correlates is crucial for elucidating the mechanisms underlying neurodevelopmental and psychiatric disorders.

Recent studies have begun leveraging animal-to-animal differences to probe how the brain generates diverse behaviors^6–8,11,13^. For example, one study found a large variability among rats in their responses to unfamiliar sounds during an auditory decision-making task, and this variability led to mechanistic insights into how neural circuits resolve uncertainty^7^. Nevertheless, most previous investigations were single-laboratory studies with small sample sizes, inherently limiting statistical power and constraining computational analyses of the origins of variability. Consequently, we still lack a clear understanding of how animal-to-animal variability is structured and how it is manifested in brain-wide neural activity. Furthermore, a scalable, data-driven method for characterizing inter-animal variability across large cohorts is still missing.

Here, we address these challenges by leveraging data from the International Brain Laboratory (IBL), in which more than 100 C57BL/6 mice were trained on an identical visual decision-making task under fully standardized conditions^5^, and brain-wide activity was recorded with Neuropixels probes^14^. In this task, a mouse rotates a wheel counter-clockwise or clockwise depending on whether a grating appears on the right or left side of a monitor, respectively. The experiment was conducted across twelve laboratories, yet housing, training, and equipment were standardized to minimize extraneous variability. Behavioral performance and block-structure dependence were consistent across labs, although the number of sessions required to reach proficiency differed slightly^5,15.^ Likewise, Neuropixels recordings from identical brain sites showed similar neuronal yields, firing rates, and LFP power across laboratories, with modest lab-to-lab differences in task-related encoding^16^. This small across-laboratory variability makes the IBL dataset ideally suited for studying animal-to-animal variability, because the remaining differences are likely to reflect fundamental biological sources rather than experimental artifacts. However, inter-animal variability in the IBL data has not yet been systematically examined. Moreover, prior behavioral analyses have focused primarily on psychometric performance^5,15,17,18,^ leaving chronometric aspects of task engagement, where richer behavioral repertoires and stronger inter-animal differences may reside, largely unexplored.

In this work we begin by analyzing the distribution of reaction times (RTs) across animals and identify two interesting features. First, mice frequently make extremely fast responses by initiating wheel movement right before stimulus onset. Second, they occasionally respond very slowly, such that RTs form a bimodal distribution on a logarithmic scale, with a secondary peak around 3 second. Analyzing their behavioral characteristics, we show that early and late RTs likely correspond to anticipatory and inattentive responses, respectively. Although most mice display both behaviors, the proportion varies widely: some animals predominantly make early responses, whereas others favor late responses. To capture this difference we introduce an anticipatory-tendency index, defined as the difference between the fractions of early and late responses. This index is stable across sessions, and female mice exhibit a slightly but significantly higher anticipatory tendency relative to males.

To investigate inter-animal variability more deeply, we develop a data-driven method that learns a low-dimensional animal embedding jointly with a deep neural network that predicts choice and RT. The learned embeddings improve prediction in held out sessions and reveal a salient structure: the primary embedding axis captures individual differences in RT, while the secondary axis reflects left/right biases of animals.

Finally, we link behavioral variability to neural dynamics by analyzing brain-wide recordings. We hypothesized that inter-animal differences arise from variations in the depth of the underlying neural landscape, which predict systematic differences in autocorrelation timescales. We tested this prediction by measuring autocorrelation during post-task passive periods and inter-trial intervals (ITIs), using a bootstrap method to obtain unbiased estimates from short ITIs. Across both periods, the autocorrelation timescale in medial visual areas is negatively correlated with an animal’s anticipatory-tendency index. Moreover, the cortex-wide relative timescales exhibits the same negative correlation, supporting our hypothesis.

## RESULTS

### Fast anticipatory and very slow inattentive features in the reaction time distribution

The IBL experiment is a decision-making task for mice. In each trial, a grating stimulus appears on either the left or right side of a screen positioned in front of a head-fixed mouse (Fig. 1A). The task requires the mouse to turn a wheel clockwise if the stimulus appears on the left, and counter-clockwise if it appears on the right. For brevity, we refer to the correct movement in response to a stimulus on the right (left) side as a right (left) motion. If the mouse performs the task correctly, it receives a small volume of sucrose water as a reward; otherwise, a burst of noise is delivered. If no choice is made within 60 seconds, the trial times out. The next trial begins automatically after a short interval (approximately 1 second) and a quiescent period (0.4–0.7 seconds). If the mouse moves the wheel more than 2^°^ during the quiescent period, the period resets, and a new quiescent interval must be completed before the stimulus appears (see Supplementary Fig. S1A for the trial timeline). Stimulus contrast is randomly selected from 100%, 25%, 12.5%, 6%, or 0%. An auditory cue (5 kHz tone) signals the onset of the stimulus, allowing the mouse to detect the start of a trial even in low contrast trials. The probability of the stimulus appearing on the left or right is modulated in a block-wise fashion. In the initial ∼100 trials of a session, stimulus locations are unbiased (50% left, 50% right), after which they become biased towards one side (80:20). Block changes occur every 20–100 trials and are not signaled to the animal.

**Figure 1:**
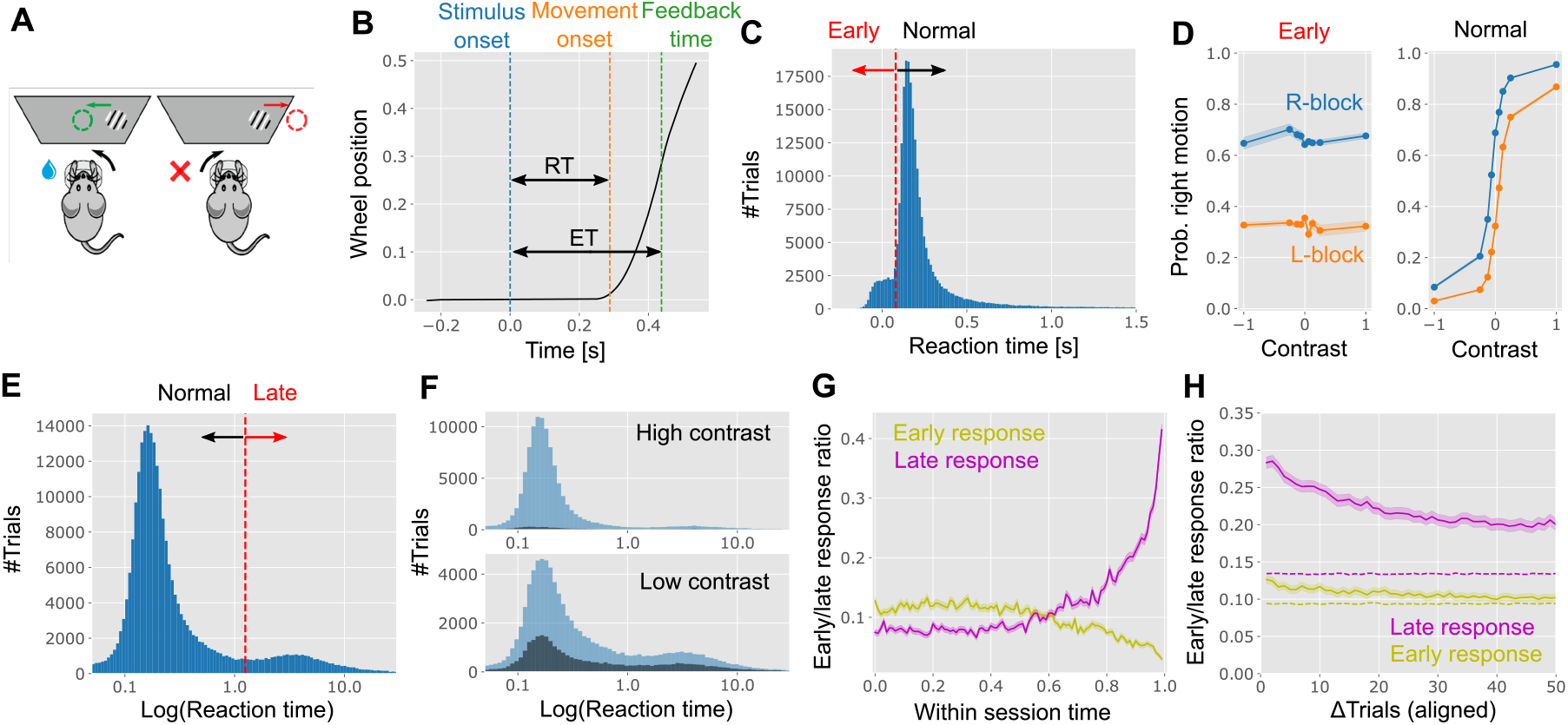
Task description and RT statistics. **(A)** Schematic of the task. **(B)** Definition of RT and ET. We defined the movement onset time as the point where a mouse started to make a significant wheel movement (orange line), and RT as the duration between the movement onset time and the stimulus onset time. Feedback time is the time point where the trial ends and the mouse receives either a water reward or a short wait time (green line). ET was defined as the duration between the feedback time and the stimulus onset time. **(C)** RT distribution across all animals and trials (n = 267,474 trials). **(D)** Psychometric curves of the wheel movement direction under early (RT < 0.08s) and normal (0.08s ≤ t < 1.25s) RTs. The contrast dependence of the probability of right motion in early response trials was not statistically significant (L-block: F-stat= 1.1, *p* = 0.36, ANOVA; R-block: F-stat= 1.4, *p* = 0.14, ANOVA). **(E)** RT plotted in a logarithmic scale. Note that bins are taken on a logarithmic scale, meaning that bins for a longer RT contain exponentially longer intervals. **(F)** The same as panel E, but divided into high (c = 25% and 100%) and low (c = 0%, 6%, and 12.5%) contrasts. Dark blue regions represent incorrect trials whereas light blue regions represent correct trials. **(G)** The ratio of early and late responses as a function of the relative position of a trial in a session. Early and late responses were defined as RT < 0.08 s and 1.25 s ≤ RT. **(H)** The early response probabilities at trial *t* + ΔTrials given an early response at trial *t* (yellow), and the corresponding curve for the late response probability (magenta). Dashed lines represent the baseline estimated from shuffling (magenta line converges to the baseline level around Δ*Trial* ≈ 500). Shaded area in panel D represents the standard error of the means (SEM) over trials, whereas the shaded areas in panels G and H are SEM over sessions.

Approximately half of the mice learned to perform the task with no significant performance differences across laboratories^5^. However, task performance as measured by psychometric curves does not fully capture the complexity of behavior^19,20^. In particular, reaction time (RT) may reveal additional behavioral variability across animals.

We examined RT distributions in expert mice, defined as those that met the criteria for electrophysiological recordings^5^. We defined the first movement onset time as the point when the mouse begins a significant wheel movement (orange vertical dotted line in Fig. 1B; see *Reaction time analysis* in Methods). RT was then defined as the interval between stimulus onset (blue line) and the first movement onset. A trial ends when the mouse moves the wheel 35 degrees in either direction, at which point it receives a reward or a noise. We refer to this endpoint as the feedback time (green line), and the duration from stimulus onset to feedback time as the execution time (ET). We analyzed RTs from 267,474 trials across 415 sessions by 132 mice with electrophysiology recordings. These sessions were chosen to study behavior of expert animals and to allow exploration of neural correlates in later analyses. We found two notable features in the RT distribution(Figs. 1C and 1E).

First, mice exhibited very fast, and often negative, RTs in approximately 10% of trials, indicating that some movements were initiated before stimulus onset (see Supp. Figs. S1B and S1C for example trajectories). In each trial, there is a 400–700 ms quiescent period during which the mouse must keep the wheel still. If the wheel is moved more than 2 degrees during this period, the quiescent period recurs (Supp. Figs. S1A). However, mice can circumvent this reset by making subtle movements or initiating movement just before stimulus onset, resulting in valid but negative RTs. These early responses were consistently observed across labs (yellow points in Supp. Fig. S2A).

To further investigate, we classified trials with RTs < 0.08 seconds as early response trials (red line in Fig. 1C), based on a clear inflection point in the RT distribution. Note that it is effectively impossible for a stimulus-driven response to occur within 80 ms of onset. As expected, in trials with normal RTs, choice behavior depended on both stimulus contrast and block structure (Fig. 1D, right). In contrast, in early RT trials, the probability of rightward movement was independent of stimulus side and contrast but strongly modulated by the block structure (Fig. 1D, left). This lack of stimulus dependence suggests that mice are making decisions before the stimulus is presented. Moreover, the strong influence of block structure implies that behavior is not random, but guided by their predictions about likely stimulus location. Note that we determined movement direction from the initial wheel movement, not the final choice, as mice sometimes changed their mind after an early movement (see Supp. Fig. S1C). These change-of-mind events were more frequent in early RT trials, particularly when the initial movement was incorrect (Supp. Fig. S1F; see Supp. Fig. S1G for the block dependence), supporting the idea that early responses reflect predictive behavior. Additionally, early responses were slightly more common in the first half of a session (yellow line in Fig. 1G), likely due to higher motivation from water deprivation. The ITI distribution for early RT trials was also more long-tailed than for normal RTs (Supp. Fig. S1H), indicating frequent quiescent period resets. Plotting the probability of observing an early response at the (*t* + *i*)-th trial given that the *t*-th trial was an early response, we found that the early response ratio is only slightly higher immediately following an early response trial (yellow solid vs. dashed lines in Fig. 1H; the dashed line represents the shuffled data; see Supp. Fig. S1L for interval distribution), indicating that early responses are not significantly clustered within a session. These findings overall suggest that early responses are anticipatory actions exploiting task structure.

Second, we observed a subset of late RT trials (Fig. 1E; see Supp. Figs. S1D and S1E for example trajectories). When plotting positive RTs on a log scale, we found a bimodal distribution, with a secondary peak around 3 seconds. We defined late response trials as those belonging to this second peak, using a threshold of 1.25 seconds (red dashed line in Fig. 1E). Late responses occurred predominantly during low-contrast trials (contrast ≤ 12.5%; Fig. 1F) and were associated with higher error rates, approaching chance level under a low contrast (Supp. Figs. S1I and S1J), suggesting disengagement.

Although standard drift-diffusion models with urgency signals predict rising error rates with increased RT^21,22^, they do not explain the long-tailed RT distribution observed here. It is more likely that mice were inattentive during these trials, rather than deliberating longer. Prior analysis of the same dataset showed larger pupil sizes before long RT trials^23^, consistent with shifts in engagement^24,25^. Similarly, GLM-HMM analyses have linked disengagement states to long RTs^17^.

Late response trials were more frequent toward the end of sessions (purple line in Fig. 1G), likely due to satiation. Due to this concentration toward the end of a session, the late response ratio was about 15% higher following late response trials (solid vs. dashed magenta lines in Fig. 1H). To control for this effect, we excluded the final 40 trials of each session in all subsequent analyses (see further analyses on the satiation effect in the next section), except for the animal embedding analysis, where within-session effects were explicitly modeled. Overall, the results above indicate that late responses primarily reflect disengagement or inattention.

### Animal-to-animal variability in fast-and-slow responses

Our analysis of the RT distribution revealed two notable behavioral patterns: early, anticipatory responses and late, disengaged responses. This raised the question of whether all mice exhibit both types of responses, or if these behaviors are specific to certain individuals. To investigate this, we computed the early and late response ratios for each animal, defined as the fraction of trials with early or late RTs, respectively, out of all trials (Fig. 2A). Notably, some animals showed a strong bias toward early responses, while others were more prone to late responses, revealing substantial inter-animal variability. To quantify this bias, we defined an Anticipatory Tendency Index (ATI) as the difference between an animal’s early and late response ratios (see Methods). A positive ATI indicates a tendency toward anticipatory (early) responses (yellow points in Fig. 2A), whereas a negative ATI suggests a predominance of late, disengaged responses (purple points).

**Figure 2:**
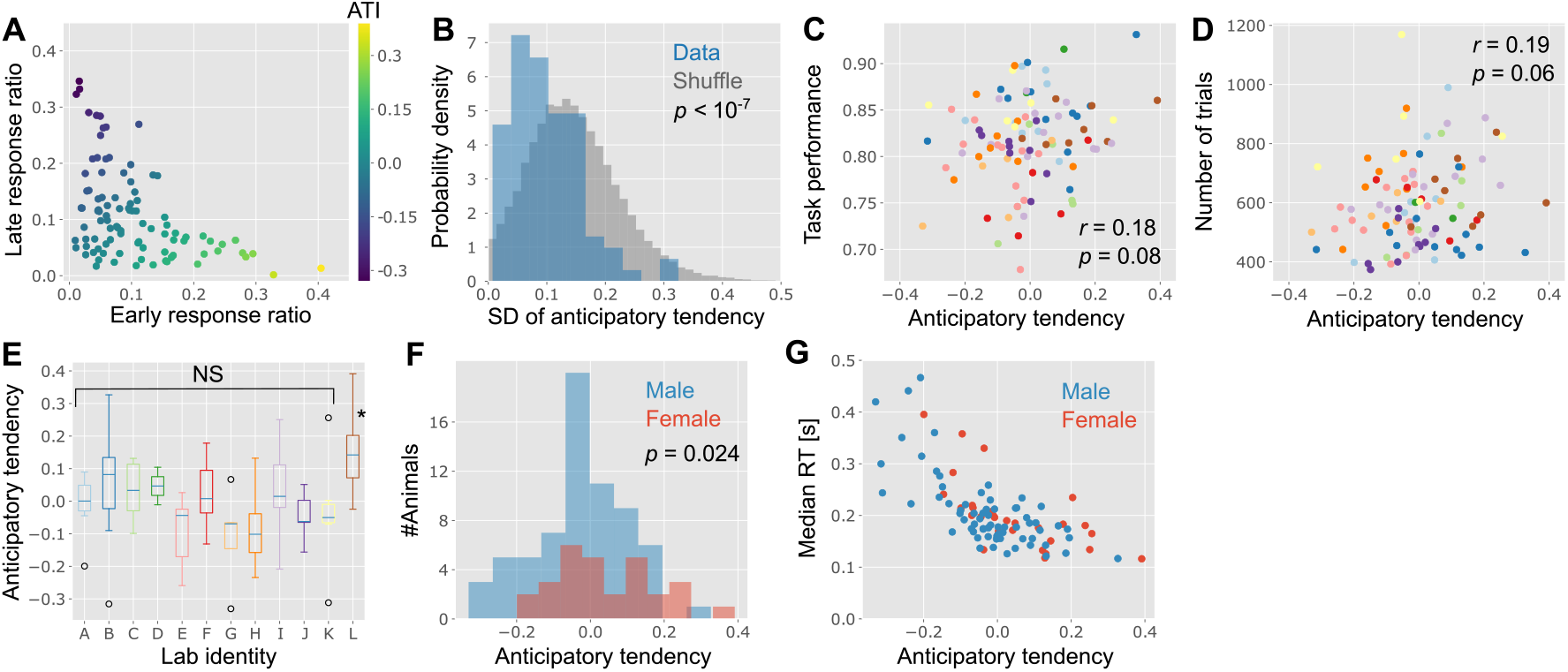
Individual variability in anticipatory tendency of movement initiation. **(A)** Mouse-to-mouse variability in early/late response ratio (*N* = 96 animals). Each point represents one mouse, and points are color-coded by the value of the anticipatory tendency index. **(B)** The standard deviation (SD) of the anticipatory tendency over sessions calculated for each mouse, compared to random shuffle (*D*(96, 96000) = 0.3, *p <* 10^*−*7^; KS-test (two-sided)). In shuffling, we randomly attributed sessions to each animal and estimated the across-session variability for each animal. **(C, D)** The anticipatory tendency is not significantly correlated with the average task performance (C; Pearson’s *r* = 0.18, *p* = 0.08) or the number of attempted trials per session (D; *r* = 0.19, *p* = 0.06) across mice. Here, mice are color-coded by the lab identity specified in panel E. **(E)** Relationship between the anticipatory tendency and lab identity (*F* (11, 84) = 3.01, *p* = 0.002; ANOVA over all labs. *F* (10, 77) = 1.90, *p* = 0.058 ANOVA over labs A-K). **(F)** Sex dependence of the anticipatory tendency (*t*(49.6) = 2.33, *p* = 0.024; Welch’s t-test on population means (two-sided); *N*_male_ = 67, *N*_female_ = 29). **(G)** The relationship between the anticipatory tendency and median RT across mice.

Importantly, ATI values were generally consistent across sessions within the same animal. We assessed this by computing the standard deviation of ATI across sessions for each mouse. Most animals showed lower session-to-session variability than would be expected by chance, based on shuffled session data (Fig. 2B). ATI was not strongly correlated with overall task performance (Fig. 2C), total number of trials performed in a session (Fig. 2D), the number of sessions required to reach expert-level performance (Supp. Fig. S2B), or animal ages (Supp. Fig. S2E).

Mice with a broad range of ATI values were observed in nearly all laboratories, although one lab (Lab L) exhibited a notably higher fraction of animals with high ATIs (*p* = 0.058, one-way F-test excluding Lab L; Fig. 2E). Interestingly, female mice showed slightly higher ATI values than males, though the effect size was modest (two-sided Welch’s t-test, *p* = 0.024; *N*_male_ = 67, *N*_female_ = 29; Fig. 2F). This finding aligns with a recent study reporting faster average reaction times in female mice compared to males^4^.

Because we found a significant shift in the ratio of early and late responses (Fig. 1G), variability in the timing of session termination could, in principle, contribute to the observed across-animal variability. To assess this possibility, in Supp. Figs. S2G-H, we plotted the within-session change in the late response ratio for different sexes and labs, respectively. Notably, although the baseline late response ratio during the first 60-70% of the session differed between male and female mice, and lab L showed a lower late response ratio in this period (thick brown line in Supp. Fig. S2H), the increase in the late response ratio toward the end of the session was similar across sexes and labs. Moreover, we observed qualitatively similar across-animal variability even when we excluded the last 200 trials (instead of the last 40 trials; Supp. Fig. S3). These results suggest that fatigue/satiation affects mice in a broadly comparable manner, and that the observed across-animal differences are unlikely to be explained by variability in the timing of session termination.

Reaction speeds, when measured using the median RT, were also consistent within individual animals across sessions (Supp. Fig. S2C). However, differences in median RT were relatively small among animals with a positive ATI (Fig. 2G). This supports the notion that ATI more effectively captures meaningful inter-animal variability in response patterns than simple metrics like median RT.

### Animal embedding analysis

The results above indicate the presence of individual variability along the anticipatory-inattention axis. This raises the question of whether this axis is the dominant factor underlying animal-to-animal variability, or whether other dimensions also contribute substantially. Moreover, it remains unclear if the observed variability improves behavioral predictions.

To explore these questions, we conducted a data-driven analysis of the variability using a deep neural network. Specifically, we built a feedforward neural network to predict the choice, reaction time (RT), and execution time (ET) of each trial, given the stimulus contrast, block, trial number (i.e., its relative position within a session), and animal ID (Fig. 3A; see *Animal embedding algorithm* in Methods). Conceptually similar to a Generalized Linear Model for behavioral prediction, the model incorporates trainable animal embedding components that capture features that vary across individuals. It also uses nonlinear transformations to learn complex interactions among input variables. The animal ID is initially encoded as a 95-dimensional one-hot vector (corresponding to the 95 animals), which is then linearly projected into a four-dimensional animal embedding space (green layer in Fig. 3A). By training the network to optimize prediction performance, the embedding space learns low-dimensional representations that capture latent factors contributing to individual variability in choice, RT, and ET.

**Figure 3:**
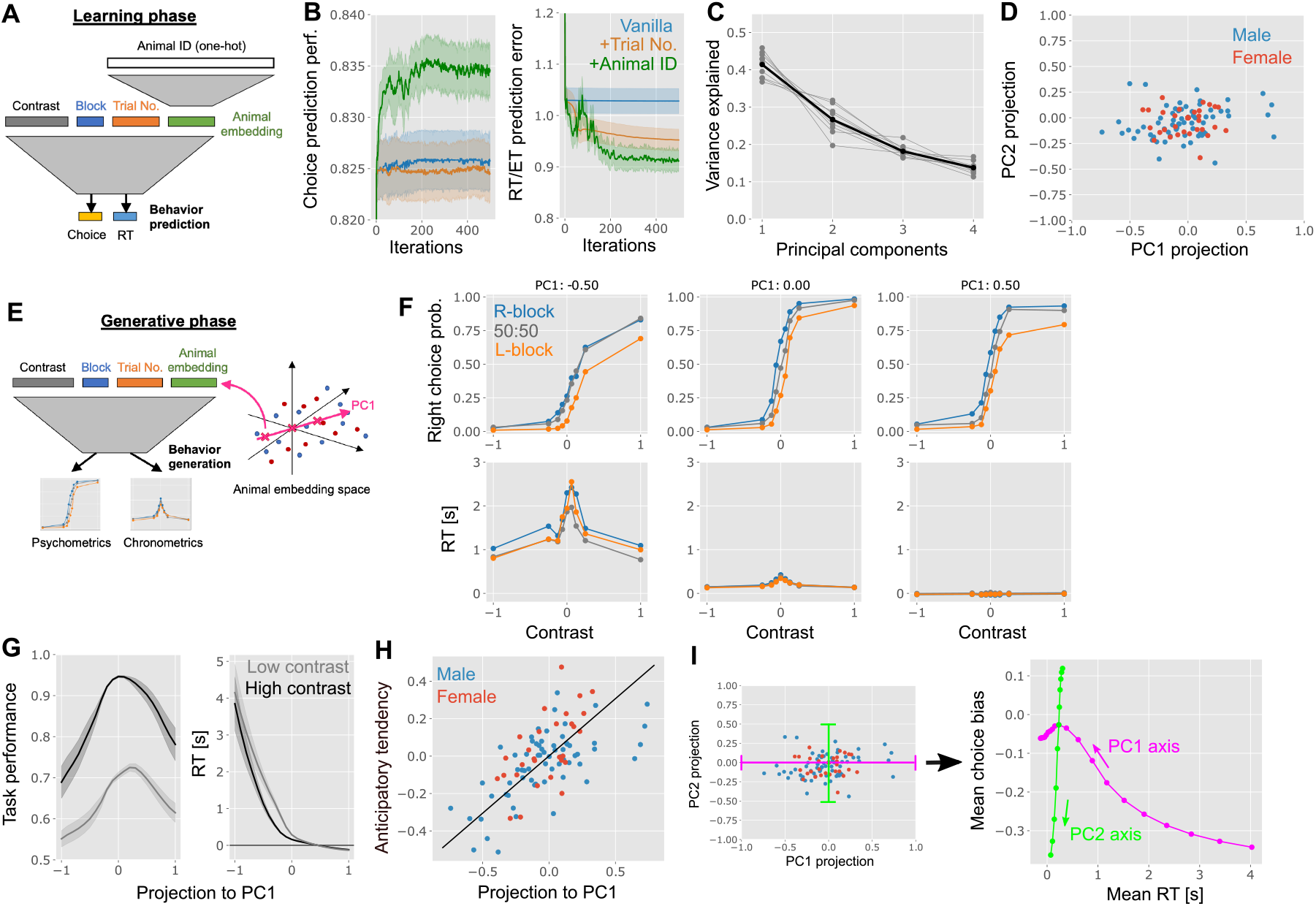
Data-driven analysis of individual variability via animal embedding. **(A)** Schematic figure of the network we trained for predicting the choice and reaction/execution time. See *Animal embedding algorithm* in Methods for the details. **(B)** Choice and RT/ET prediction performance in held-out session. The vanilla network only uses the stimulus contrast and block for predictions, while the ‘+Trial No.’ model also uses the trial number, and the ‘+Animal ID’ model incorporates animal ID for prediction. Error bars are the standard errors over 10 random training/test splits. **(C)** Normalized eigen-spectrum of the learned animal embedding. Thin lines represent results from 10 simulations with random train/test splits. **(D)** Projection of the embedding vector of each animal onto the space spanned by the two principal directions of the animal embedding matrix. **(E)** Schematic of the ideal animal psychometric and chronometric curves generation. **(F)** Expected psychometric and chronometric curves along the PC1 axis of the animal embedding. **(G)** Task performance and RT along the PC1 axis averaged over contrasts and blocks. Shaded areas represent the standard error of the mean over fittings from 10 random seeds. **(H)** Projection of animal ID to PC1 of the animal embedding space correlates with the anticipatory tendency index (*r* = 0.65, *p <* 10^*−*3^, *N* = 95). **(I)** Projection of PC1 and PC2 axes of the animal embedding (pink and green lines in the left panel) onto the space spanned by the mean RT and the mean choice bias (curves in the right panel).

We trained this model on 334 sessions from 95 expert mice and evaluated its performance on held-out sessions. Sessions were split semi-randomly to ensure that each session in the test dataset had at least one corresponding session from the same animal in the training dataset. After training, the model incorporating animal ID outperformed the model without it in predicting both choices and RTs on held-out sessions (green vs. blue lines in Fig. 3B). Moreover, when we shuffled the animal ID, the performance on both choice and RTs prediction also went down significantly (Supp. Fig. S5C). These results highlight consistent behavior across sessions within individual animals and indicates that the model captures features enabling cross-session prediction. Additionally, the trial’s relative position within a session was important for predicting RT (orange vs. blue line in Fig. 3B, right), consistent with the observed within-session shift in late-response rates (Fig. 1G).

To further investigate the structure of individual variability, we analyzed the learned animal embeddings. We performed eigen-decomposition on the weight matrix that maps one-hot animal IDs into the low-dimensional embedding space. We found that the first principal component (PC1) accounted for over 40% of the variance captured in the embedding space (Fig. 3C). When projecting all animals into the PC1–PC2 space, we observed a continuous distribution without distinct clusters (Fig. 3D).

A key advantage of our embedding approach is that it enables behavioral simulation of a hypothetical animal, allowing us to interpret the embedding axes. We aligned the embedding vector along the PC1 axis and used it to estimate average choice, RT, and ET across different stimuli and blocks, yielding psychometric and chronometric curves (Fig. 3E). Note that, because we fitted log-transformed RT (see Methods), point-estimate of RT generated from the learned model captures the central tendency of RT more reliably than the mean of raw RT. Along the PC1 direction, we observed a drastic shift in chronometric curves: RTs exceeded one second for animals with negative PC1 projections (bottom-left panel, Fig. 3F) and became shorter or even negative for animals with positive projections (bottom-right). The psychometric curve showed a pronounced leftward bias in the high-RT region (top-left), a pattern observed in some mice. When averaging chronometric curves to estimate mean RT, we found that RT decreased monotonically, but nonlinearly, along the PC1 direction (right panel, Fig. 3G). Task performance also varied systematically: deviations in either direction along PC1 reduced accuracy (left panel, Fig. 3G). This suggests that both impulsivity and inattention are associated with poorer task performance, relative to a more balanced behavioral state. This relationship was less apparent when analyzing performance as a function of anticipatory tendency alone (Fig. 2C), possibly because higher-order components obscure this trend. Notably, projection onto PC1 was highly correlated with ATI defined earlier (Fig. 3H), suggesting that the anticipatory-inattentive axis is a major axis of individual variability.

In contrast, PC2 primarily captured variations in psychometric bias (left vs. right) with only minor effects on RT (Supp. Fig. S5B). When plotting average choice bias and RT across PC1 and PC2 axes, we observed nonlinear but approximately orthogonal trends (Fig. 3I). This suggests that individual variability is at least two-dimensional: although highly inattentive mice tend to exhibit left bias, animals across a broad range of anticipatory-inattentive profiles can display left, right, or no bias at all.

To compare this observed structure in the individual variability with previously reported individual variability in a similar experimental protocol^6^, we next examined the slope difference (Supp. Fig. S4). Following Liebana et al. (2025)^6^, we defined the right (left) slope as the absolute difference between the probability of right movement under high contrast (25% and 100%) and zero contrast, and then defined the slope difference as the difference between the two slopes. While this slope difference was correlated with PC1 projection (Supp. Fig S4D; *r* = −0.23, *p* = 0.02), the correlation with PC2 projection was stronger (Supp. Fig S4E; *r* = 0.44, *p* = 10^*−*5^), implying that the slope difference mainly captures bias rather than decision strategy in the IBL data, potentially due to differences in the task and training protocols.

### A neural landscape hypothesis on individual variability

The behavioral analyses above reveal a low-dimensional structure underlying animal-to-animal variability, with a principal axis corresponding to the anticipatory-inattentive tendency. In the following sections, we explore the potential neural origins of this variability. One plausible hypothesis is that differences in the neural landscape underlie the observed behavioral differences, which has previously been proposed in the context of autism spectrum disorders^26^, excitatory/inhibitory imbalance^27^, and sex differences in bandit task behaviors^4^. Figure 4A illustrates this conceptual model.

**Figure 4:**
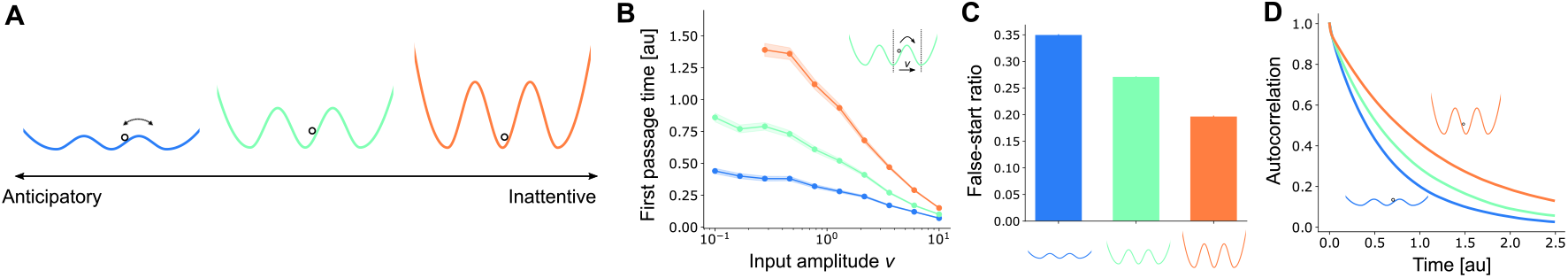
A neural landscape model on individual variability. **(A)** Schematic of the model. **(B)** First passage time (the duration it takes for the neural state to move from one to another) under three neural landscapes as a function of the input amplitude. We initiated the state at the center and estimated the time until it traverses to the right attractor as illustrated in the inset. **(C)** False-start ratio under the three models. Here, we defined the false-start as the transition to another state before the input is provided to the model. **(D)** autocorrelation of dynamics under the three models. See *Neural landscape model* in Methods for details.

Neural population dynamics in the brain are often described as evolving within a landscape that contains attractor states, activity patterns corresponding to specific behavioral states, actions, or sensory inputs. In a shallow neural landscape (blue curve in Fig. 4A), the system can more easily transition between attractors. For example, if one attractor corresponds to a “hold” state and another to a “rightward movement preparation” state, then spontaneous transitions across the states may result in premature responses or false starts, as seen in anticipatory animals. In contrast, a deep landscape stabilizes neural states and reduces transitions, but may impair sensitivity to weak stimuli, potentially contributing to inattentiveness.

To test this idea, we implemented a simple dynamical model simulating state transitions under varying landscape depths. When constant input was applied, we found that the first-passage time, the time it takes for the network to transition between states, is longer in the deep landscape (Fig. 4B; see *Neural landscape model* in Methods). This effect was negligible when inputs were strong enough to reliably drive transitions (right side of Fig. 4B), but became noticeable under weak input conditions. Moreover, the false-start ratio, defined as the proportion of trials in which state transitions occurred before the input stimulus, was higher in the shallow landscape (Fig. 4C).

This toy model thus offers a mechanistic hypothesis for the neural basis of inter-animal variability. It also leads to a testable prediction: namely, that animals with shallow neural landscapes (i.e., anticipatory mice) should exhibit shorter autocorrelation timescales in their neural dynamics, while those with deeper landscapes (i.e., inattentive mice) should show longer timescales. This follows from the idea that shallow landscapes allow rapid transitions, causing autocorrelation to decay quickly, whereas deep landscapes promote state persistence and thus slower decay (blue vs. orange curves in Fig. 4D). We therefore hypothesize that autocorrelation timescales in neural activity will systematically differ between anticipatory and inattentive animals.

The neural landscape model formulated above assumes the presence of task-relevant attractor states. Although this assumption is partially supported by prior analyses of the same dataset^14,28^, it remains unclear whether task-relevant neural activity states are similar across trials with diverse RTs. We thus visualized average neural activity trajectory during trials in example sessions using Gaussian Process Factor Analysis^29,30^ (GPFA), which extracts low-dimensional latent dynamics from population activity (see *Neural trajectory and representational similarity analyses* in Methods). Lines in Figs. 5A and 5B represent the mean neural activity trajectory in the inferred latent space for five RT bins from two example sessions (A: prelimbic area (PL); B: anterior visual area (VISa)). We averaged the trajectory for left and right trials separately and excluded incorrect or zero-contrast trials. In both areas, neural dynamics were similar across RT bins, though the inferred trajectories differed across regions and sessions. The pre-stimulus points cluster across RT groups, consistent with a local neural state associated with the pre-stimulus period (gray cloud in Fig. 5A left). Similarly, activity around stimulus onset and around movement/feedback forms separable clusters (blue and green clouds, respectively), consistent with additional locally defined task-related states.

**Figure 5:**
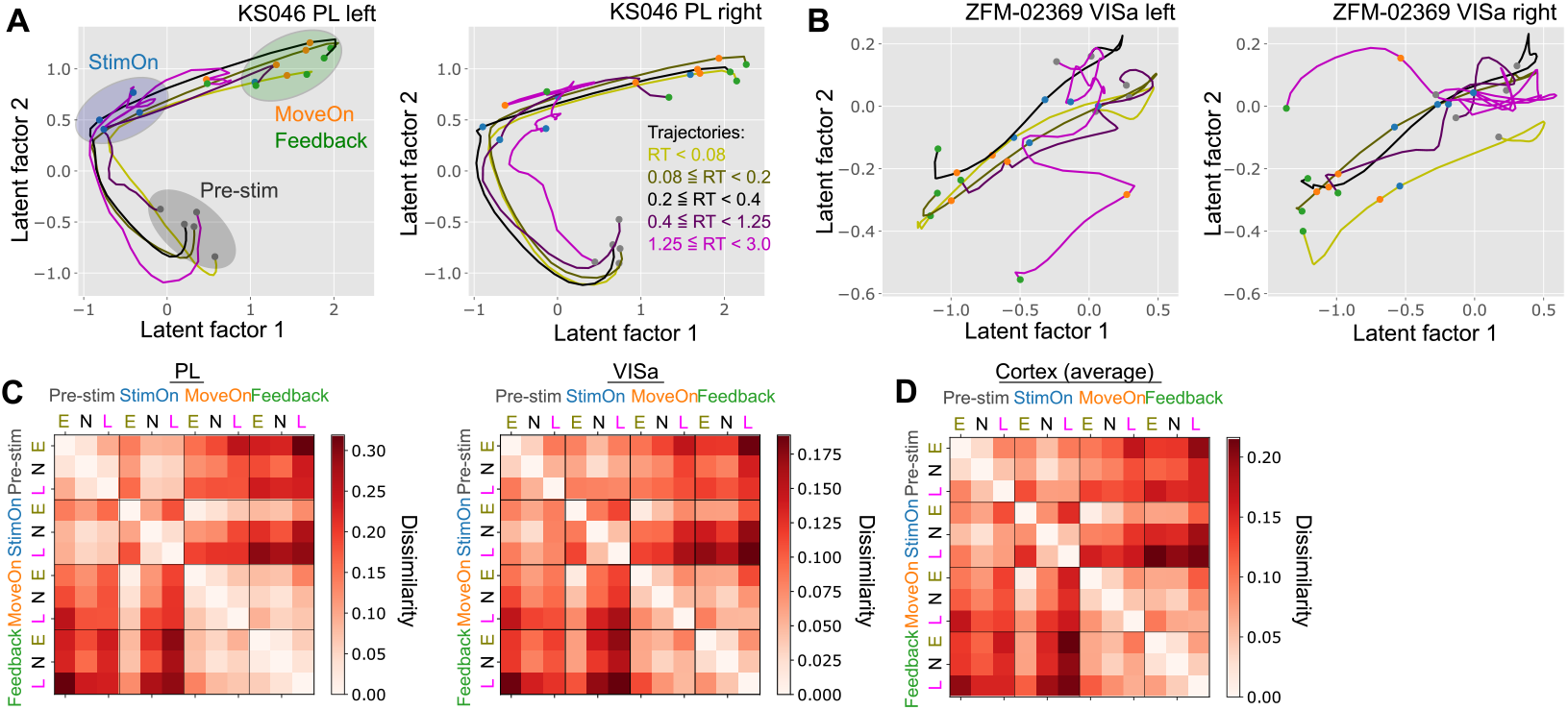
Illustration of state transitions in local population activity. **(A, B)** Example within-trial neural trajectories from two recordings (A: prelimbic (PL) area from animal KS046; B: anterior visual area (VISa) from animal ZFM-02369). Only correct, non-zero-contrast trials were included, and left- and right-stimulus/movement trials are plotted separately (but fitted jointly). The trajectories correspond to trial-averaged activity in reaction-time (RT) bins (*RT <* 0.08, 0.08 ≤ *RT <* 0.2, 0.2 ≤ *RT <* 0.4, 0.4 ≤ *RT <* 1.25, and 1.25 ≤ *RT <* 3.0). Each trajectory begins at pre-stimulus activity (gray points) and then evolves through stimulus onset (blue), movement onset (orange), and feedback (green). **(C)** Representation dissimilarity matrices for PL (left) and VISa (right). E, N, and L denote early-, normal-, and late-RT trial groups, respectively defined as: early: *RT <* 0.08, normal: 0.08 ≤ *RT <* 1.25, and late: 1.25 ≤ *RT*, as before. Larger dissimilarity values indicate greater differences in population activity between conditions (see Methods). **(D)** Same as panel C, but averaged across all recorded cortical regions.

Figures 5C and 5D show the representational dissimilarity^31,32^ across different task periods (pre-stimulus/stimulus onset/movement onset/feedback periods) and RT groups (Early/Normal/Late), measured by maximum mean discrepancy (MMD) between population activity distribution^31,32^ (C: PL and VISa; D: cortex-wide; see Methods). On average, neural activity in each task period was similar across RT groups. In particular, the population activity during the pre-stimulus period of an early-RT trial was much closer to the activity during the pre-stimulus period of a normal-RT trial than to the feedback period activity of early-RT trials (top-left vs top-right squares in Figs. 5C and 5D). One exception is the stimulus-onset period activity of early-RT trials, which resembled movement onset activity of other RT groups than their stimulus onset activity (‘StimOn E’ columns and rows in Figs. 5C and 5D). These results indicate the presence of task-relevant states not depending strongly on RT, implying the presence of stable neural states. However, we also observed a large variability in extracted neural trajectories across sessions and regions. We thus focus on the autocorrelation analysis below, which is robust against the variability in specific task representation across regions and sessions.

### Comparison with autocorrelation during passive period

To test the predictions derived from the neural landscape model, we analyzed the neural activity of each animal across the brain recorded with Neuropixel probes 14 and compared it to their anticipatory tendency. We focused specifically on the passive period—the time following the task during which no external stimulus was presented (Fig. 6A) to minimize the influence of task-related behav-iors on neural activity.

**Figure 6:**
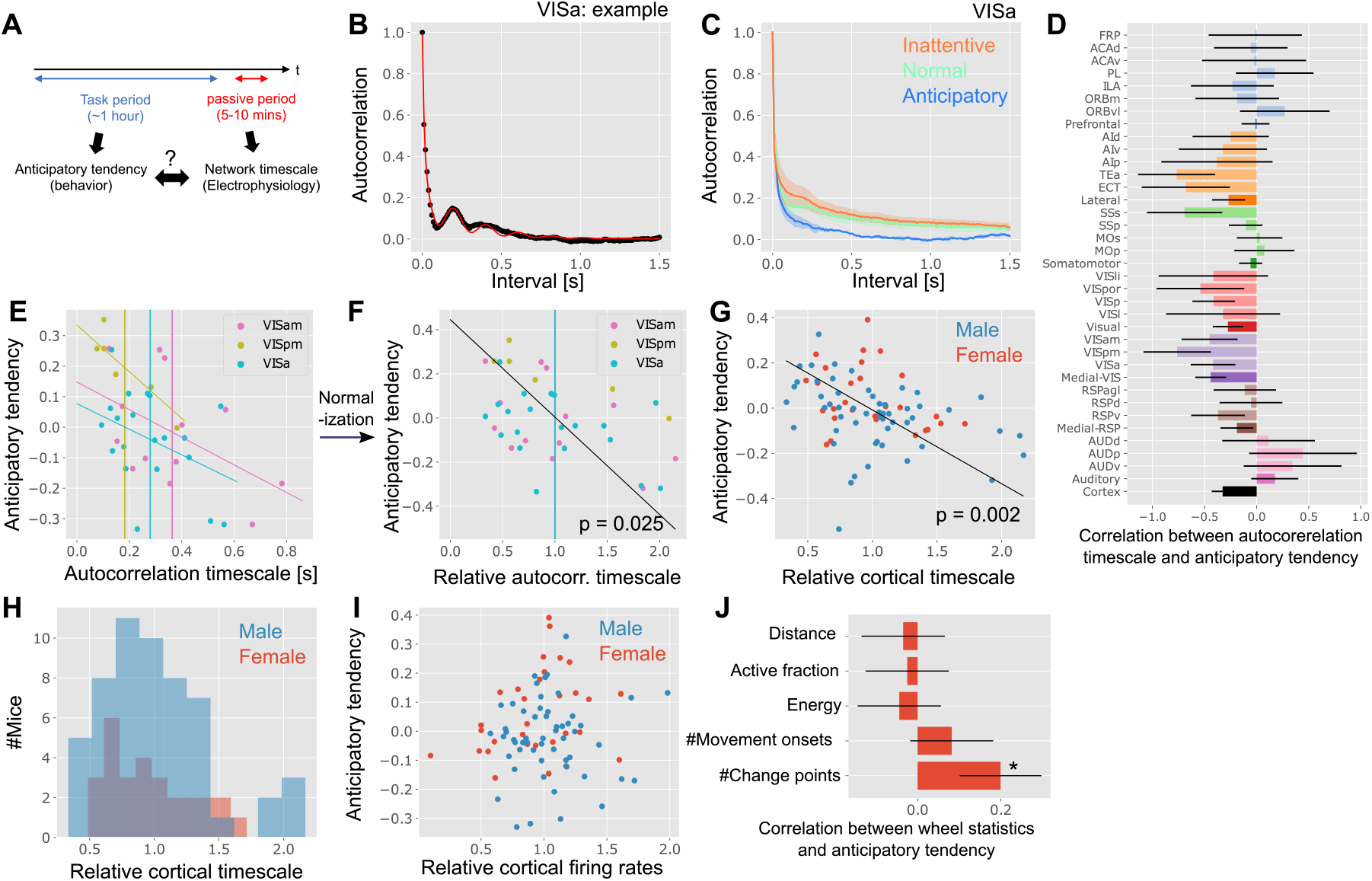
Comparing the individual variability in anticipatory tendency and autocorrelation timescales during passive period. **(A)** Schematic of the autocorrelation analysis. **(B)** An example curve of the autocorrelation of population activity in one recording from the VISa region (black points) and its fitting (red curve). **(C)** Average autocorrelation of VISa population activity for inattentive (ATI < -0.15), normal, and anticipatory (ATI > 0.15) mice. Error bars represent the standard error of the mean across sessions. **(D)** Correlation between the anticipatory tendency and autocorrelation timescale of various regions (pale bars) and divisions (dark bars). The acronyms follow the Allen Brain Atlas^33^. The error bars represent standard errors without Bonferroni correction. **(E**,**F)** Estimated autocorrelation timescales and their relationship with session-wise anticipatory tendency for VISam (pink), VISpm (yellow), and VISa (blue) regions. Each point represents one session. In panel F, we divided the autocorrelation timescales by the region-wise mean timescales (vertical lines in panel E). In panel E, statistics were (*r, p*) = (− 0.45, 0.09), (− 0.76, 0.003), (− 0.42, 0.05) for VISam, VISpm, and VISa, respectively (permutation-test). In F, we had *r* = − 0.44, *p* = 0.025, *N* = 39. **(G)** Correlation between the anticipatory tendency and the relative cortical timescale (*r* = − 0.33, *p* = 0.002, *N* = 83). Each point represents one mouse. **(H)** Histogram of the relative cortical timescale of male and female mice (*p* = 0.71, permutation test; *N*_male_ = 55, *N*_female_ = 28). **(I)** Relative cortical firing rate did not show correlation with the anticipatory tendency (*r* = 0.08, *p* = 0.50). **(J)** Correlation between wheel statistics and anticipatory tendency. The number of change points exhibited a statistically significant correlation with the anticipatory tendency (*r* = 0.23, *p* = 0.018; permutation test), while the other wheel metrics did not.

We estimated autocorrelation at the population level by calculating the population firing rate within each brain region. This approach allowed us to infer autocorrelation from the relatively short passive periods reliably. Because autocorrelation during this period often exhibited oscillatory components, we fitted a double-exponential function with an oscillatory term to the autocorrelation curves (Fig. 6B; see *Autocorrelation timescale estimation* in Methods); this improved the fitting consistently across areas (Supp. Fig. S6A). We defined the autocorrelation timescale as the longer of the two time constants from the fitted double-exponential terms. The estimated timescales were broadly consistent with those reported in previous work^34^, although we did not observe a clear correlation with the cortical hierarchy scores reported in prior studies^35,36^ (Supp. Fig. S6C). We excluded sessions with poor fit quality or timescale estimates that were clear outliers (see Methods for exclusion criteria).

Figure 6D shows the correlation between the autocorrelation timescale of 29 cortical regions and each animal’s anticipatory tendency during the preceding task. We excluded regions with fewer than five valid sessions. While many cortical areas exhibited negative correlations between autocorrelation timescale and anticipatory tendency, the variance was high. We thus grouped regions into functional divisions based on prior anatomical classifications^35^, normalizing each session’s estimated timescale by the average for that region (Figs. 6E and 6F). We split the medial division into visual and retrosplenial (RSP) areas, given their distinct roles and the potential importance of medial visual areas in this task^16^.

With this grouping, we found significant negative correlations between session-wise anticipatory tendency index (ATI) and autocorrelation timescales in the medial visual areas (*p* = 0.025, Bonferroni-corrected across 7 divisions). Moreover, the average autocorrelation curves estimated for VISa resembled the model prediction (Fig. 6C vs. 4D). This result aligns with our model’s prediction: anticipatory animals exhibit faster neural dynamics (i.e., shorter timescales) compared to inattentive animals. We next computed a relative cortical timescale for each animal by averaging the normalized autocorrelation timescales across all recorded regions. Since Neuropixels recordings were performed acutely and the recording site varied daily for each animal^14^, we aggregated data across days. For example, if VISam was recorded on Day 1 and MOp on Day 2, the animal’s relative cortical timescale was taken as the mean of these two sessions’ normalized timescales (see Methods). We found that the relative cortical timescale showed a significant negative correlation with ATI values across animals (*p* = 0.002, Pearson’s r = −0.33, *N* = 83 animals; Fig. 6G), again supporting the neural landscape hypothesis. Although animals with longer cortical timescales were often male, the effect of sex on timescale was not statistically significant (*p* = 0.71, permutation test; Fig. 6H).

We also examined the relationship between relative cortical firing rates and anticipatory tendency. Using the same normalization and aggregation procedure, we found no clear correlation between firing rate and ATI (Fig. 6I), although relative firing rates of auditory areas was correlated with session-wise ATI (Supp. Fig. S6D).

Finally, because mice often engaged with the wheel even during passive periods, we examined various wheel movement statistics as proxies for spontaneous behavior. Total wheel movement distance, fraction of time spent moving, wheel energy, and the number of movement onsets did not show significant correlations with ATI (Fig. 6J). However, we did find a weak but statistically significant correlation between ATI and the number of trajectory change points in wheel movement (*p* = 0.04; Fig. 6J and Supp. Fig. S6H). This suggests that anticipatory animals may be somewhat more fidgety even during the passive period.

### Comparison with autocorrelation during inter-trial interval (ITI)

The results above suggest that animals with a higher anticipatory tendency exhibit faster neural dynamics during the passive period, consistent with predictions from the neural landscape hypothesis. To further test this relationship, we next analyzed neural activity during the inter-trial interval (ITI), as illustrated in Fig. 7A.

**Figure 7:**
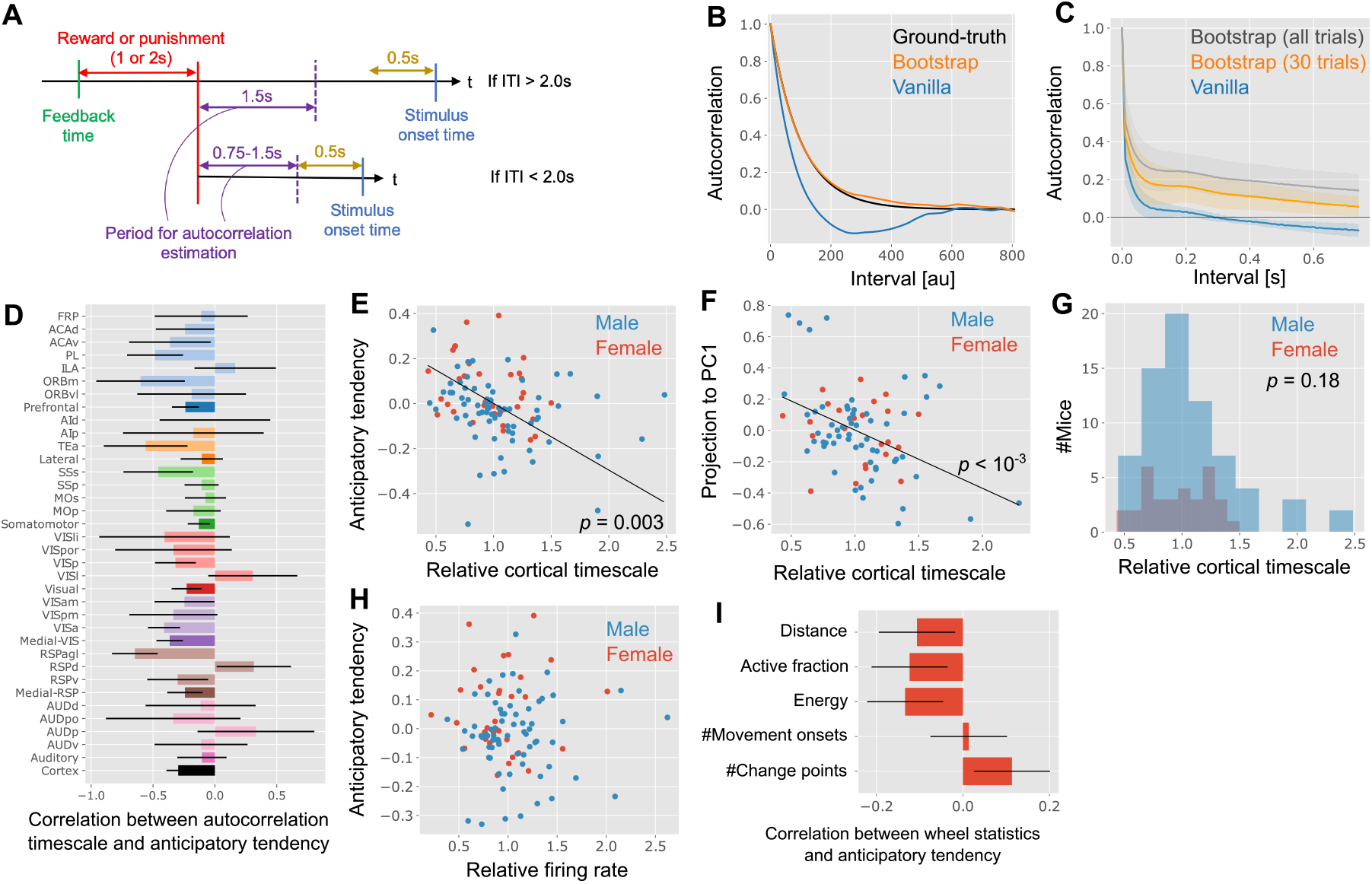
Comparing the individual variability in anticipatory tendency and autocorrelation timescales during ITI. **(A)** Schematic of the ITI and the selection of period for the autocorrelation estimation. **(B)** Autocorrelation estimation with vanilla or bootstrap methods. Here, the ground truth was generated artificially by sampling sequences from a Gaussian Process with an exponential decay kernel function. **(C)** The average autocorrelation curve of VISa population activity estimated with vanilla and bootstrap methods. The error bars are the standard error of the means over recordings. **(D)** Correlation between the anticipatory tendency and autocorrelation timescale of various regions (pale bars) and divisions (dark bars). The error bars are the standard errors without Bonferroni correction. **(E, F)** Correlation between the relative cortical timescale and the anticipatory tendency (E; *r* = − 0.3, *p* = 0.003, *N* = 103; permutation-test) and the projection to PC1 of the animal embedding (F; *r* = − 0.37, *p <* 10^*−*3^, *N* = 82). Each point represents one mouse. **(G)** Histogram of the relative cortical timescale (*p* = 0.18, permutation-test; *N*_male_ = 70, *N*_female_ = 33). **(H)** Relative cortical firing rate did not show clear correlation with the anticipatory tendency (*r* = − 0.02, *p* = 0.85). **(I)** Correlation between wheel statistics and anticipatory tendency. Anticipatory tendency did not show statistically significant correlation with the wheel statistics listed above.

Analyzing neural dynamics during ITIs presents two key technical challenges. First, estimating autocorrelation from short-duration ITIs is non-trivial. Standard autocorrelation methods become biased when the interval length is comparable to the underlying timescale, due to intrinsic correlations between neural activity and its trial-wise mean^37–39^. Figure 7B illustrates the discrepancy between the ground-truth autocorrelation and a naive (“vanilla”) estimation method, even when averaging across thousands of ITIs (black vs. blue lines; see *Autocorrelation timescale estimation* in Methods).

To address this, we developed a simple bootstrapping method in which the mean population activity is estimated using nearby ITIs rather than on a trial-by-trial basis. This enables computationally efficient estimation of autocorrelation curves and offers a practical alternative to more complex generative model-based methods^39,40^. We prove that this method yields asymptotically unbiased autocorrelation estimates when ITIs are independently drawn from a Gaussian process (see *Supplementary Text* for proof). Applied to synthetic data generated from a Gaussian process, the bootstrap method closely recovered the ground-truth autocorrelation (orange vs. black lines in Fig. 7B).

Applying this approach to neural recordings during ITIs, we observed that the vanilla method produced autocorrelation curves with a negative dip (blue line, Fig. 7C), as expected. In contrast, the bootstrap method, with a 30-trial window for mean and variance estimation, produced smooth, naturalistic autocorrelation decay curves (orange line, Fig. 7C). Using very large trial windows (e.g., all trials) introduced constant positive bias due to violation of the i.i.d. assumption. We fixed the trial window size at 30 for our analyses, though comparable results were obtained with 10- and 50-trial windows (Supp. Figs. S8C and S8D).

A second technical concern is that anticipatory mice tend to exhibit longer ITIs than inattentive ones (Supp. Fig. S2D), which could confound autocorrelation estimates. To control for this, we restricted analysis to the first 1.5 seconds of each ITI (Fig. 7A). For ITIs shorter than 2 seconds, we analyzed the period from ITI onset to 0.5 seconds before the stimulus onset, to reduce contamination from motor preparation.

Using the bootstrap method, we estimated autocorrelation timescales for each recording and assessed their correlation with ATI values at both the session and animal levels. Consistent with the passive period findings, we observed a significant negative correlation between relative autocorrelation timescale in medial visual areas and session-wise ATI (Fig. 7D; *p* = 0.008, Bonferroni-corrected across divisions; see also Supp. Fig. S7A for VISa-specific results). Furthermore, when averaging across all recorded regions per animal, the resulting relative cortical timescale also showed a significant negative correlation with animal-wise ATI (Fig. 7E; Pearson’s r = −0.3, *p* = 0.003, *N* = 103 animals). In addition, this timescale was significantly correlated with each animal’s projection onto PC1 of the animal embedding space (Fig. 7F). We confirmed that these results were robust to different neuron inclusion criteria and ITI definitions (Supp. Fig. S8).

As in the passive period analysis, sex differences in cortical timescales were not statistically significant (Fig. 7G), and cortex-wide firing rates did not correlate with anticipatory tendency (Fig. 7H, Supp. Fig. S7D). We also found no strong behavioral confounds: wheel movement metrics during ITIs, including movement distance, energy, active fraction, number of movement onsets and change points, showed no significant correlation with anticipatory tendency (Fig. 7I, Supp. Figs. S7F and S7G).

To further examine the robustness of our analysis, we next conducted the auto-correlation analysis using single-neuron-wise fitting rather than population-level fitting. Even with this highly stochastic neuron-wise estimation of autocorrelation and its timescale, we still observed a robust negative correlation between the estimated relative cortical timescale and the anticipatory tendency of animals (Supp. Figs. 9E–G). Moreover, the estimated autocorrelation timescale scaled with cortical hierarchy (Supp. Fig. 9H), consistent with a recent analysis of the same data^41^. Nevertheless, these correlations were not evident when we applied the same analysis to the passive-period data (Supp. Figs. 9B and D), except for the medial visual areas (Supp. Fig. 9C), potentially due to high stochasticity and oscillatory activity (Supp. Fig. 9A and Fig. 6B).

This robust variability in ITI autocorrelation led us to ask whether an animal’s anticipatory tendency also influences the encoding of prior (i.e., block) information, which has been shown to be decodable across a wide range of brain regions^28^. Following Findling et al. (2025)^28^, we quantified prior decoding performance in each region using the coefficient of determination (*R*^2^) (Supp. Fig. S10A). Consistent with previous work, in normal-RT trials the Bayesian-optimal prior could be robustly decoded from pre-stimulus activity in many cortical regions (Supp. Fig. S10B; example shown for MOs). In contrast, decoding accuracy estimates for early- and late-RT trials were highly variable, likely due to the smaller number of trials in these subsets. At the animal level, subjects with poor prior decoding tended to show negative anticipatory tendency, but this relationship was only marginal overall (Supp. Fig. S10C; *r* = 0.24, *p* = 0.051; see legend for details).

Overall, the consistency between passive-period and ITI analyses reinforces our main conclusion: animals with higher anticipatory tendencies exhibit faster cortical dynamics, especially in medial visual areas, providing evidence for a neural correlate of individual behavioral variability.

## DISCUSSION

In this study, we investigated behavioral variability among mice and its neural correlates in a decision-making task. By analyzing reaction times across more than quarter million trials from over 100 animals, we identified distinct properties of both negative-RT and very slow RT trials, trial types that are usually discarded in standard RT analyses, and uncovered substantial inter-animal variability in the proportion of these trials. Using an animal-embedding model, we further discovered a low-dimensional representation of this variability that predicts each animal’s behavior. Finally, autocorrelation analysis of cortex-wide activity during passive and ITI periods revealed robust neural correlates of anticipatory tendency in autocorrelation timescale. Together, these findings support the hypothesis that differences in the “depth” of the landscape of cortical dynamics may underpin individual behavioral variability.

Our work also includes two methodological contributions. The first is the animal embedding algorithm (Fig. 3). While the use of deep neural networks is common in large-scale behavioral data analysis^42^, previous studies either pooled data from all animals or trained a separate network for each individual animal. In contrast, our proposed approach captures inter-animal variability in a way that enables interpretable predictions of behavior. The second methodological contribution is an efficient algorithm for estimating autocorrelation from a short ITI period. We demonstrate that our method provides an unbiased estimate of autocorrelation even with limited data, unlike naive estimation approaches. This method also enables faster estimation than previous approaches based on generative model estimation^39^. However, it assumes that ITIs are independently sampled, an assumption that is expected to be violated over a long session (Fig. 7C).

Latent trajectories we found are consistent with region-wise, task-relevant neural landscapes (Fig. 5), but do not provide direct evidence of their existence. Additionally, the systematic differences in autocorrelation timescales align with predictions derived from across-animal variability in the neural landscape (Figs. 6 and 7), while not fully ruling out potential confounding factors. Although we focused on passive and ITI periods to minimize task-related confounds, the observed negative relationship between anticipatory bias and cortical time-scale may still have behavioral origins. Most animals moved the wheel significantly during fewer than 5 % of passive periods (Supp. Fig. S6G), yet we detected a weak positive correlation between the number of wheel-movement change points and anticipatory tendency (Supp. Fig. S6H). This suggests that highly anticipatory animals may fidget more even during passive periods, modulating brain-wide dynamics^43^. We also observed that anticipatory mice exhibit a more heavy-tailed ITI distribution than inattentive mice (Suppl. Fig. S2D). Although our analyses focused on the initial segment of each ITI to mitigate this effect, this observation indicates cross-animal differences in mental states during ITIs. Moreover, the between-animal differences in cortical timescale we report may not reflect intrinsic circuit properties; Neuromodulatory signals are known to reshape network autocorrelation^44,45^, raising the possibility that animals with different ATI spend different amounts of duration in distinct brain states.

While the medial visual cortex consistently showed the strongest neural–behavioral coupling, this may partly reflect the denser sampling of VISa and VISam, which were targeted for reproducibility assessments^16^. We indeed obtained a high yield from VISa (Supp. Fig. S6J). Although we also obtained high yields from primary visual cortex (VISp) and primary somatosensory cortex (SSp), these regions are large and functionally heterogeneous, which may introduce greater session-to-session variability.

Because all subjects were C57BL/6J mice, the variability we observed is unlikely to have a genetic origin. Prior studies have shown that behavioral variability can be shaped by multiple factors^10^, including the sex and age of the animals^1,4,46^, the physical characteristics of the vivarium and laboratory^47,48^, in-cage social hierarchy^2,3^, and implementation details of behavioral tasks^49,50^. Sex explained a modest fraction of the variance in our analysis, but the neural signature of this effect was not apparent. Housing conditions present another plausible source: the mice were group-caged whenever possible, and social hierarchy within cages can induce behavioral divergence even in genetically identical animals^2,3^. In particular, non-dominant mice tend to display more anxiety-like behavior^2^, which could influence their task engagement.

Structural and functional brain signatures of behavioral variability across individuals have been extensively studied in humans^51–53^, but a circuit-level understanding requires studies in animal models. In rodents, Kurikawa et al. reported behavioral variability in response to unfamiliar auditory cues and linked it to individual differences in neural landscapes; however, their analysis was limited by low statistical power due to a small sample size^7^. Chen et al. identified sex differences in strategy during a bandit task in mice and attributed them to differences in the depth of explore and exploit states^4^, but they did not examine the underlying neural correlates. Neural correlates of variability in contextual decision-making strategies^8^ and learning trajectories^6^ have also been explored recently. Nevertheless, systematic, data-driven accounts of behavioral variability and its brain-wide correlates remain lacking. Our work addresses this gap by linking individual variability in cortical dynamics to behavioral variability along the anticipatory–inattentive axis.

Several recent studies have analyzed behavior and neural activity in the IBL task^5,14–18,28,54^. Notably, fluctuations in task engagement within single sessions have been reported^17,18^, and concurrent work by Laranjeira et al. has identified multiple session-wise behavioral phenotypes^55^. Our results complement these findings by revealing cross-animal variability in task engagement and linking it to cortical neural dynamics.

Finally, our observations resonates with human resting-state fMRI studies reporting altered neural landscape dynamics in autism spectrum conditions^26,56,57^. In particular, Watanabe et al. recently showed that autistic individuals exhibit more rigid network dynamics, resembling the inattentive landscape in our model^57^. Nevertheless, it should be noted that the local region-wise neural landscape we analyzed here is distinct from the global neural landscape studied in those works. The sex difference we observe in anticipatory bias may likewise relate to the male predominance in autism^58^. Applying our framework to mouse models of autism-associated mutations therefore represents an important avenue for future research.

## MATERIALS AND METHODS

### Reaction time analysis

#### Definition of reaction time (RT)

For each trial, we defined the movement onset time as the first time point where the speed of the wheel exceeds 0.5 [rad/s] after 300 ms before the stimulus onset time and following more than 50 ms of stationary wheel duration during which speed is below 0.5. We picked 300 ms prior to the stimulus onset as the earliest movement onset of a trial because we found a large number of trials where a mouse starts moving the wheel right before the stimulus onset. RT was defined as the duration between the movement onset time and the stimulus onset time. If the movement onset occurs before the stimulus onset, it takes a negative value, otherwise, RT takes a positive value. Execution time (ET) was defined as the duration between the feedback time (the time when reward or noise feedback is provided) and the stimulus onset time. See Supp. Fig. S1A for the trial timeline.

#### Anticipatory tendency index

We defined the anticipatory tendency index by:

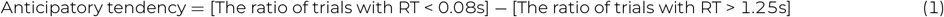

We excluded the last 40 trials from the estimation of the anticipatory tendency and other behavioral variables, to minimize the effect of satiation. A small number of trials (2, 387 trials out of 267, 474 total trials) did not have a valid RT either due to timeout without movement or potential mechanistic error. We did not exclude these trials from the denominator, as the fraction is small (<1%), and the fraction of invalid trials was not correlated with the anticipatory tendency. Unless otherwise stated, we estimated the ratio for each animal by concatenating trials from all the sessions. In Figs. 6C-6E and 7D (excluding the last ‘cortex’ bars in 6E and 7D), we estimated the anticipatory tendency in a session-wise manner for a comparison with session-wise autocorrelation timescale.

#### Exclusion criteria

In our analysis, we only included sessions with electrophysiological recording to minimize the variability originating from task learning. Additionally, we only included sessions with more than 400 trials (exclusive). This is because a session is terminated at the 400th trial if the animal appears to be clearly disengaged 5. In Fig. 2, to achieve a robust estimation of the animal-wise anticipatory tendency and other behavioral variables, we only included animals with two or more valid recording sessions (*N* = 96 animals).

### Animal embedding algorithm

#### Datasets

We used behavioral data from recording sessions to study the behavioral variability among animals that have already learned the tasks. As before, we only included the sessions that had more than 400 trials (exclusive). For this embedding analysis, we defined a valid trial where task variables are not NaN, and RT and ET satisfy, 0.03*s* ≤ *RT* − 0.2 and 0.03*s* ≤ *ET*. The latter two conditions were introduced to make the logarithmic values of RT and ET valid and stable. We excluded one animal that had sessions where the number of valid trials was less than 80 %. We only included animals with two or more electrophysiology sessions as before. These criteria resulted in 371 sessions from 95 animals. We split the sessions into training and test sessions with a 90:10 split. We made sure that for each test session, there is at least one session from the same animal in the training dataset, but the training-test split was done randomly otherwise.

#### Inputs and outputs

Input vectors were constructed as a composition of the representation of the stimulus contrast, task block, trial number, and animal ID. Stimulus contrast vector is a nine-dimensional one-hot vector corresponding to one of nine stimulus contrasts (-1.0, -0.25, -0.125, -0.06, 0.0, 0.06, 0.125, 0.25, 1.0). Here, negative and positive contrasts represent stimuli on the left and right sides. Similarly, a block vector is a three-dimensional one-hot vector representing one of three task blocks (0.2, 0.5, 0.8). For the representation of the relative position of a given trial in a session, we used a Gaussian receptive field model. We set the activity of the *i*-th position encoding neuron given the *t*-th task as exp 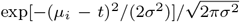, where *µ*_*i*_ is the selectivity of the *i*-th neuron and *σ*^2^ is the variance. We used *N*_*pos*_ = 4 neurons for position encoding, and set 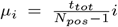 (for *i* = 0, 1, 2, 3), and 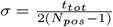.

Animal embedding was constructed by assigning a one-hot vector to all 95 animals first, then projecting this 95-dimensional vector to a four-dimensional latent space using a learnable linear weight. The full input was then constructed by concatenating vectors representing the stimulus contrast, block, relative trial position, and animal embedding, resulting in a 20-dimensional vector.

As the target outputs, we consider two objectives simultaneously: choice and time for trial initiation and execution. For the choice, we consider the standard binary classifier with two-dimensional outputs corresponding to left and right choices. For the task time prediction, we constructed two regression targets: RT and ET. Because the reaction time shows a bimodal distribution on the logarithmic scale, here we considered the prediction of time on the log-scale. To this end, we converted the reaction time and execution time to log (*RT* + 0.2) and log (*ET*). We excluded trials where these conversions are not well-defined.

#### Network architecture and learning algorithm

As depicted in Fig. 3A, we used a feedforward neural network model to predict animal choice, RT, and ET. The model architecture is written as

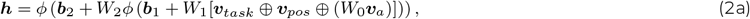

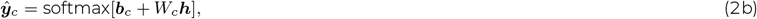

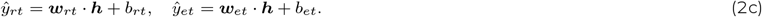

Here, *W*_0_, *W*_1_, *W*_*c*_ are synaptic weight matrices, and ***b***_1_, ***b***_2_, ***b***_*c*_, *b*_*rt*_, *b*_*et*_ are bias parameters. *ϕ* is the rectified linear function (*ϕ*(*x*) = max(0, *x*)) and ***a*** ⊕ ***b*** represents vector concatenation. We set the layer width of both hidden layers to be 32, thus the size of matrices *W*_1_, *W*_2_, and *W*_*c*_ are 32 × 20, 32 × 32, and 2 × 32, respectively. ***w***_*et*_ and ***w***_*et*_ are 32-dimensional vectors. The loss function was defined as

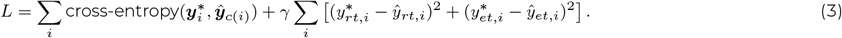

We set the balance parameter between the two losses *γ* as *γ* = 1, because the residual loss on the choice and RT/ET were roughly the same. We trained the network using Adam optimizer^59^ with learning rate 0.001 and momentum 0.9. We used each session as a mini-batch and trained the network for 500 epochs. All weights were trained in an end-to-end fashion, meaning that the learned animal embedding is locally optimized for behavioral prediction. We implemented the algorithm using the Flax library^60^.

#### Task generation and analysis

For the latent analysis, we studied the structure of 4 × 95 dimensional embedding matrix *W*_0_ using eigen-decomposition. In Figs. 3F and Supp. Figs. S5A and S5B, we generated the psychometric and chronometric curves using trained networks. In Fig. 3F, we fixed the trial number input to *t* = 400 with *ttot* = 800, and set the animal embedding vector to −0.5***v***_1_, **0**, and 0.5***v***_1_ from the left to right where ***v***_1_ is the principal eigenvector of the animal embedding. We then obtained the choice and RT predictions for all combinations of stimulus contrast and block to generate psychometric and chronometric curves. Similarly in Fig. S5A, we fixed the animal embedding vector to zero and set the trial number input to *t* = 0, 400, and 800 from the left to right while using *t*_*tot*_ = 800. This means that the left, middle, and right panels correspond to psychometric and chronometric curves at the beginning, middle, and end of a session.

In all the figures, we took the average over psychometric and chronometric curves generated from ten networks with different training/test splits and weight initialization. Because the learned animal embedding is sign invariant, we aligned the sign of PC1s across the ten obtained representations such that the negative sign corresponds to slower RT. Similarly, we aligned the sign of PC2s across ten networks such that the negative sign corresponds to the right bias.

### Neural landscape model

In Fig. 4, we constructed a toy model of neural landscape *f* (*x*) by

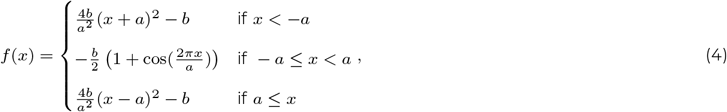

and simulated discretized dynamics by

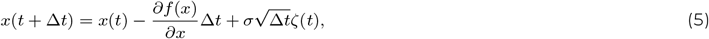

where *ζ*(*t*) is a random variable sampled from a Gaussian distribution 𝒩 (0, 1), and Δ*t* = 0.01 is the discretization bin size. We fixed *a* at *a* = 1 and changed *b* across models as *b* = 2, 4, 6. Small *b* corresponds to a shallow landscape (blue curves), while large *b* yields a deep landscape (orange curves).

In Figs. 4B and 4C, we initialized the dynamical system at *x* = 0, then ran the dynamics under *σ* = 1.75. After 50 time steps (at t = 0.5), we introduced a constant positive input *v*. We then evaluated the time it takes until the first time the state crosses the bump on the right and achieves *x > a*. This is known as the first-passage time^61^. We subtracted the first 50 time steps from the estimated duration. False-start rate was estimated by the ratio of trials where the dynamics reached *x > a* before the input is provided (i.e., within 50 time steps).

In Fig. 4D, we set the noise amplitude *σ* = 2.0, ran the dynamics for 10^6^ time steps, and estimated the autocorrelation curves.

### Autocorrelation timescale estimation

#### Autocorrelation fitting

Autocorrelation curves typically contain multiple exponential factors corresponding to neuronal and network-level autocorrelation timescales in addition to oscillatory components. We thus fitted the following function:

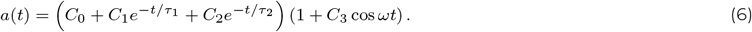

We binned population activity of a given region by 0.01-second bins, and calculated autocorrelation up to a 1.5-second delay. Parameters (*C*_0_, *C*_1_, *C*_2_, *C*_3_, *τ*_1_, *τ*_2_, *ω*) were estimated using a trust-region method implemented in the scipy curve-fit function. We sampled initial values from a uniform distribution where the ranges were chosen as [0.01, 1.0] for (*C*_0_, *C*_1_, *C*_2_, *C*_3_), [0.001, 0.03] for *τ*_1_, [0.03, 1.0] for *τ*_2_, and [0.0, 2*π/*10] for *ω*. We picked different initial ranges for *τ*_1_ and *τ*_2_ to encourage the symmetry breaking. We bounded all co-efficients to be non-negative, and *τ*_1_ and *τ*_2_ to be lower-bounded by 0.001 [second] for stability. We set the maximum iteration of curve-fit to 10^6^, and repeated fitting from 10^4^ random initializations, then picked the parameters achieving the best fit. We picked the max(*τ*_1_, *τ*_2_) as the autocorrelation timescale. This is because the exponential decay with faster timescale is expected to correspond to single-neuron level timescale, not the network timescale^39^.

In Supp Fig. 6A, we compared the fitting error of this function and a standard double exponential fitting:

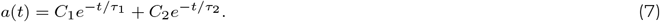

The error was defined by 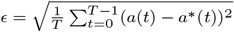, where *a*^*^(*t*) is the empirical estimate of the autocorrelation curve. In the ITI fitting, we again binned population activity of a given region by 0.01-second bin, but calculated autocorrelation only up to 0.75-second delay, as the short ITI makes estimation of a long-delay autocorrelation infeasible. We used the double exponential fitting (Eq. 7), because of this short delay and the oscillation was generally not strong during ITI compared to the passive period. We used the same fitting protocol as above, but set the maximum iteration of curve-fit to 10^5^, and repeated fitting from 10^3^ random initializations due to the presence of fewer free parameters.

#### Exclusion criteria

For the passive period analysis, we only included sessions where the passive period was clearly tagged in the data, resulting in fewer sessions than the sessions used for the ITI analysis. Moreover, we only included sessions where the total number of neurons recorded from the given region is equal to or larger than 10, and the total number of spikes in the passive period is larger than 10^4^, to reduce the noise in the empirical autocorrelation estimation. For the ITI analysis, we set the same criteria for the minimum number of neurons and the cumulative number of spikes during ITIs. We did not impose any criteria on the sorting quality for inclusion of a neuron for the analysis except for Supp. Fig. S6I and S8A.

After fitting, we imposed two more criteria: First, we only used the fitting where the fitting error is lower than 0.025 (orange vertical line in Supp. Fig. S6A). Secondly, we excluded the fitting where the estimated relative timescale is outside of 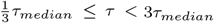, where *τ*_*median*_ is the median timescale of the given region (see black vertical line in Supp. Fig. S6B). We imposed this second criterion because it is unlikely to observe a ten-fold difference in autocorrelation timescale across individuals.

#### Unbiased autocorrelation estimation using bootstrapped mean and variance

Given a sequence 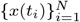 of equally sampled time points (*t*_*i*+*j*_ − *t*_*i*_ = *j*Δ*t*), the standard definition of autocorrelation follows^38,39^

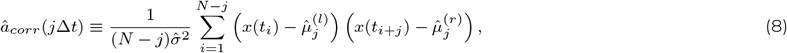

where the left and right means 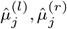 and the variance 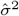 are given by

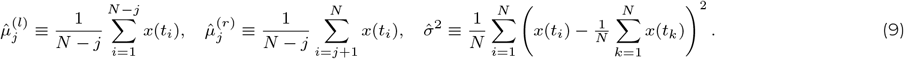

However, this estimation is known to be biased when the total duration of the sequence *t*_*N*_ − *t*_1_ is comparable to the typical auto-correlation timescale^37,38^. This is because there exists an intrinsic correlation between *x*(*t*_*i*_) and 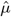 (see *Supplementary text*).

One way to overcome this bias is to consider inference of the underlying generative model^39^. However, this method is computationally expensive as it requires sample generation from the generative model at every iteration of optimization. An alternative approach we propose here is to bootstrap the mean and variance from nearby trials. More specifically, given a population activity sequence at the *k*-th trial 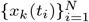, we estimate the autocorrelation at the *k*-th trial by

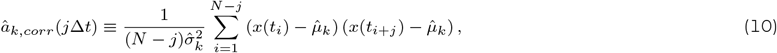

where

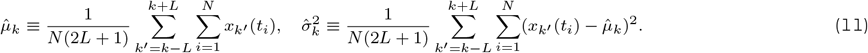

Here, *L* specifies the range of trials we use to estimate the mean and variance. Because the true mean and variance underlying ITI population activity are expected to shift during the session, it is not appropriate to use all trials for estimating 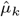 and 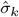. By contrast, if we only use one or two neighboring trials to estimate these values, the estimation would be unreliable. In the analysis, we fixed *L* to be 15, which yielded autocorrelation curves without a dip or a constant factor (Fig. 7C).

#### Cortex-wide relative autocorrelation timescale

We estimated the cortex-wide relative timescales plotted in Figs. 6G, 6H, and 7E-7G as follow. Let us denote the estimated autocorrelation timescale of region *r* in subject *s* by 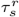, the set of recorded regions from subject *s* by Ω_*s*_, and the set of subjects with recording from region *r* by Γ_*r*_. Then the relative autocorrelation timescale of subject *s* is defined by

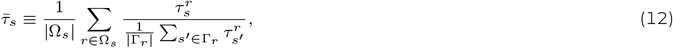

where |Ω_*s*_| and |Γ_*r*_ | represents the size of the sets Ω_*s*_ and Γ_*r*_, respectively. We used the same procedure to estimate the cortex-wide relative firing rates.

#### Neuron-wise autocorrelation timescale analysis

In the main text, to reliably estimate autocorrelation from highly stochastic neural activity over a short recording duration, we computed autocorrelation using region-wise population activity. In Supplementary Fig. S9, we additionally performed neuron-wise estimation of autocorrelation timescales. Specifically, we selected neurons with firing rates exceeding 1 Hz during the relevant period and computed spike-count autocorrelations using 0.01 s bins (see Supp. Fig. S9A for example traces). For each neuron, we fitted a double-exponential function (Eq. 7) to the autocorrelation trace. We then defined the autocorrelation timescale for each session–region pair as the mean of the longer fitted timescale across neurons recorded simultaneously in that region. We excluded neurons with poor fits (fit error > 0.01) and only included sessions where we found 10 or more neurons with a good fit for our analysis. For the passive-period analysis, we did not include an oscillatory term because oscillations were often weak in single-neuron autocorrelation curves. We additionally required QC = 1.0 as a neuron-selection criterion for the passive-period analysis. In both the passive-period and ITI analyses, all other procedures matched those used for the region-wise autocorrelation timescale estimation described above.

#### Firing rate analysis

In the firing rate analysis, we simply estimated the average firing rate per neuron during the passive period and ITI. We applied the same criteria on the minimum total number of neurons per region and the total number of spikes.

#### Wheel statistics analysis

In Figs. 6J and 7I, we calculated summary statistics of the wheel trajectories *w*(*t*) during the passive period and ITI as follow. The wheel distance was defined as the total movement of the wheel per second, 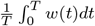. The active frac-tion is the ratio of active wheel movement period defined as 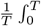 Heaviside 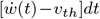 *dt* where 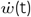 is the speed of the wheel at time *t* and *v*_*th*_ = 0.5 rad/s. The energy is given by 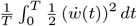. The movement onsets were calculated in the same way we calculated the movement onset of a trial (see *Definition of reaction time* in Methods), and the value was defined as the number of movement onsets per second. Lastly, the change point is the point where the velocity changes sign. Given a discrete observation of the wheel trajectory at time *t*_1_, *t*_2_, …, time point *t*_*i*_ is a change point if (*w*(*t*_*i−*1_) − *w*(*t*_*i*_)) (*w*(*t*_*i*_) − *w*(*t*_*i*+1_)) *<* 0. The number of change points was defined as the change points per second.

### Neural trajectory and representational similarity analyses

In the neural trajectory analysis, we applied GPFA^29^ to simultaneously recorded population activity in a given region. We only included correct trials with non zero-contrast stimuli where the execution time is well-defined and smaller than 10 seconds. We analyzed left and right trials jointly but plotted separately. For each trial, we set the beginning and the end of a trajectory as one second before the stimulus onset and 0.1 second after the feedback time, respectively. We then applied GPFA to the population spiking activity data using a Python library (https://github.com/aecker/gpfa). We set the number of latent variables to three, but only depicted the first two dimensions. Bin size for trajectory plotting was set to 0.05s. The mean trajectories over RT bins were estimated by taking the mean activity over trials in the latent space at each time point. For the movement onset point (orange points), the point represents the median movement onset time over trials belonging to each bin.

Representational similarity analysis was conducted in the original population activity space rather than reduced latent space. We first estimated the population activity vector ***r*** in 0.2s bins for each period. For the pre-stim period activity, we took average firing rate at 1.0-0.8 s before the stimulus onset. For stimOn, moveOn, and feedback activity, we took average over 0.2 second bins centered around the stimulus onset time, movement onset time, and feedback time, respectively (see Fig. 1B for the definitions). As in the trajectory analysis, we only included correct non-zero contrast trials with valid task execution time. We estimated the distance between population activity vectors under two conditions using maximum mean discrepancy (with diagonal components)^31,62^ using the cosine kernel, which enables us to compare two firing rate distributions, not just their means. Specifically, given a set of activity vectors under condition A, 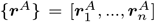, and condition B, 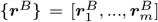, we denote the distance between {***r***^*A*^} and {***r***^*B*^} by

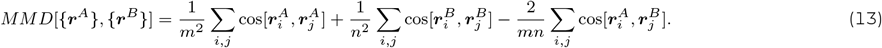

Figs. 5C and D plot this estimate for pairs of 12 conditions. For instance, the element corresponding to the (Pre-stim E, Pre-stim N) pair depicts MMD between Early-RT pre-stim period activity and Normal-RT pre-stim period activity. We only estimated the dis-tance when both condition *A* and *B* contains more than 10 trials. We took the average of the dissimilarity matrix over sessions with recording from PL (VISa) in Fig. 5C left (right). In Fig. 5D, we further took the average over all recorded cortical regions.

## DATA AVAILABILITY

Both the behavioral and neural data we analyzed are publicly available through The International Brain Laboratory.

## CODE AVAILABILITY

Codes for replicating all the results are available at https://github.com/nhiratani/IBL_variability.

## ACKNOWLEDGMENTS

The authors thank Peter Latham for his help with the project conceptualization and discussion. This work was partially supported by the McDonnell Center for Systems Neuroscience.

## SUPPLEMENTARY FIGURES

**Supplementary Figure 1:**
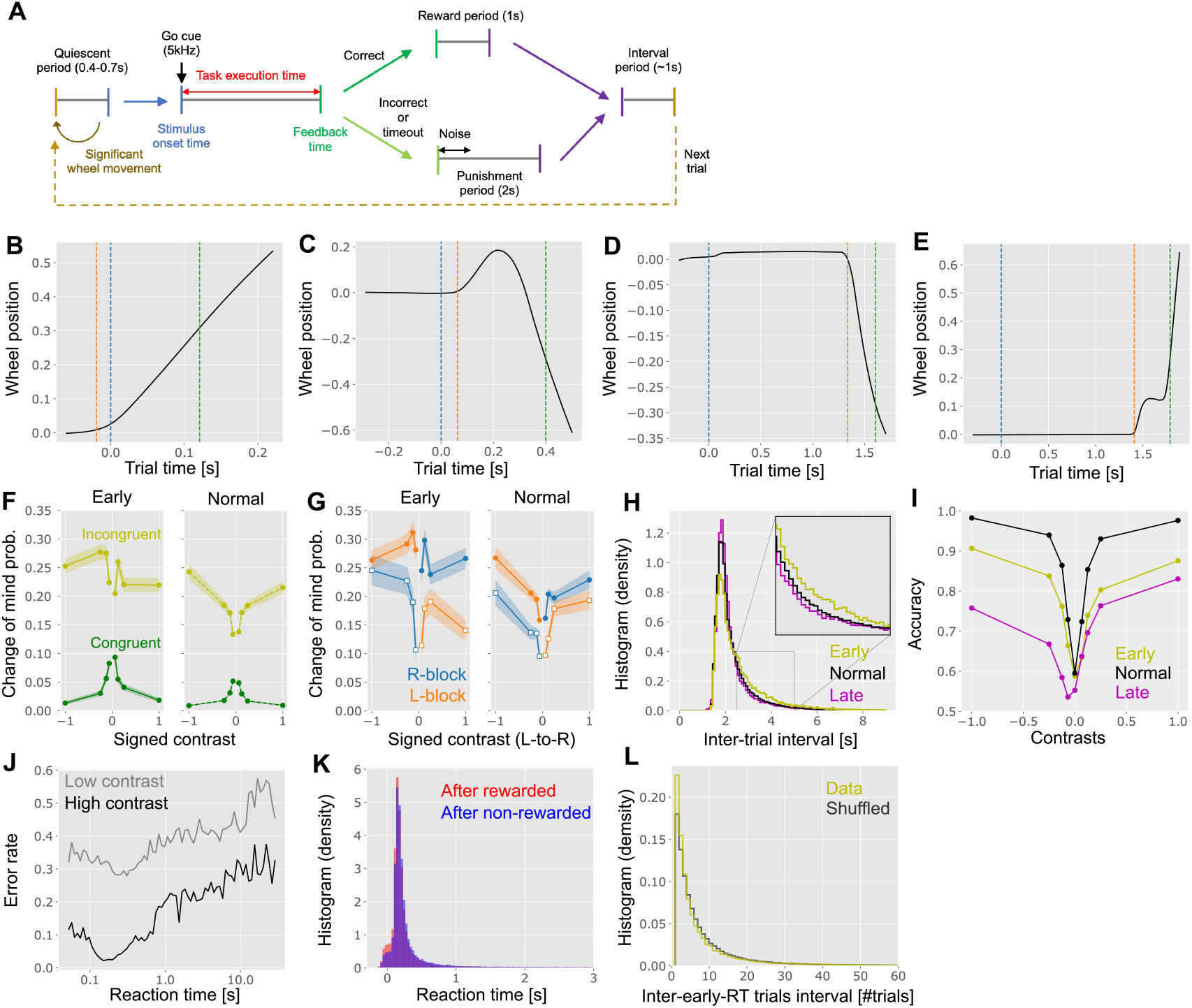
Reaction time statistics. **(A)** Step-by-step description of a trial. Note that, below, we denote the combination of the interval period and quiescent period as the inter-trial interval. **(B-E)** Examples of wheel trajectories in early response trials (B,C) and late response trials (D,E). As in Fig. 1B, blue, orange, and green lines represent the stimulus onset, movement onset, and the feedback time. While ballistic responses shown in panels B and D are the most common, there were also trials where a mouse changes their mind in the middle (panel C), or stops the wheel in the middle and moves it again in the same direction (panel E). **(F)** The probability of the change of mind in early and normal response time trials. A trial was classified as a change of mind trial if the directions of the first movement and the final choice are different as in panel C. We estimated this probability for each signed stimulus contrast in congruent and incongruent trials. Congruent trials are defined as trials where the direction of the first movement is the correct direction, whereas incongruent trials are the ones where the first wheel movement is the opposite of the correct direction. **(G)** The change-of-mind probability in incongruent trials (i.e., trials in which the initial movement direction is opposite to the correct direction), plotted separately for right and left blocks. Filled vs open markers indicate whether the stimulus side is consistent with the block prior (filled-circle: consistent; open-square: inconsistent), while the block identity is shown by color. In both early- and normal-RT trials, the change-of-mind probability is higher when the initial movement is incongruent with both the prior and the stimulus (i.e., when the stimulus is consistent with the block), compared with trials in which it is incongruent with the stimulus direction but consistent with the prior. This implies that the probability of a change of mind depends on congruence with both the stimulus direction and the block information. **(H)** Distribution of inter-trial intervals preceding early, normal, and late trials. ITI was defined as the duration between the end of the previous trial and the stimulus onset of the new trial. In a reward trial, the trial ends one second after the feedback time, while in an error trial, the trial ends two seconds after the feedback time. The inset expands the area marked by the gray box for clarity. **(I)** Performance accuracy (measured by the final movement) as a function of the stimulus contrast for the three RT groups (early/normal/late). **(J)** Error rate as a function of the reaction time in low contrast (gray) and high contrast (black) trials. **(K)** Reaction time distribution plotted separately for trials following reward and non-reward. **(L)** Histogram of the interval between two early-RT trials over all sessions. Dark gray line represents the histogram under the session-wise trial shuffling. Two histograms largely overlap with each other, indicating that the early-RT occurs in a nearly random manner, rather than having a clear clustering structure (mean and median intervals for data: (9.2, 4); for shuffled: (9.4, 4)). Note that the median interval is small because the majority of early-RT trial occurrences were concentrated in a relatively small number of sessions.

**Supplementary Figure 2:**
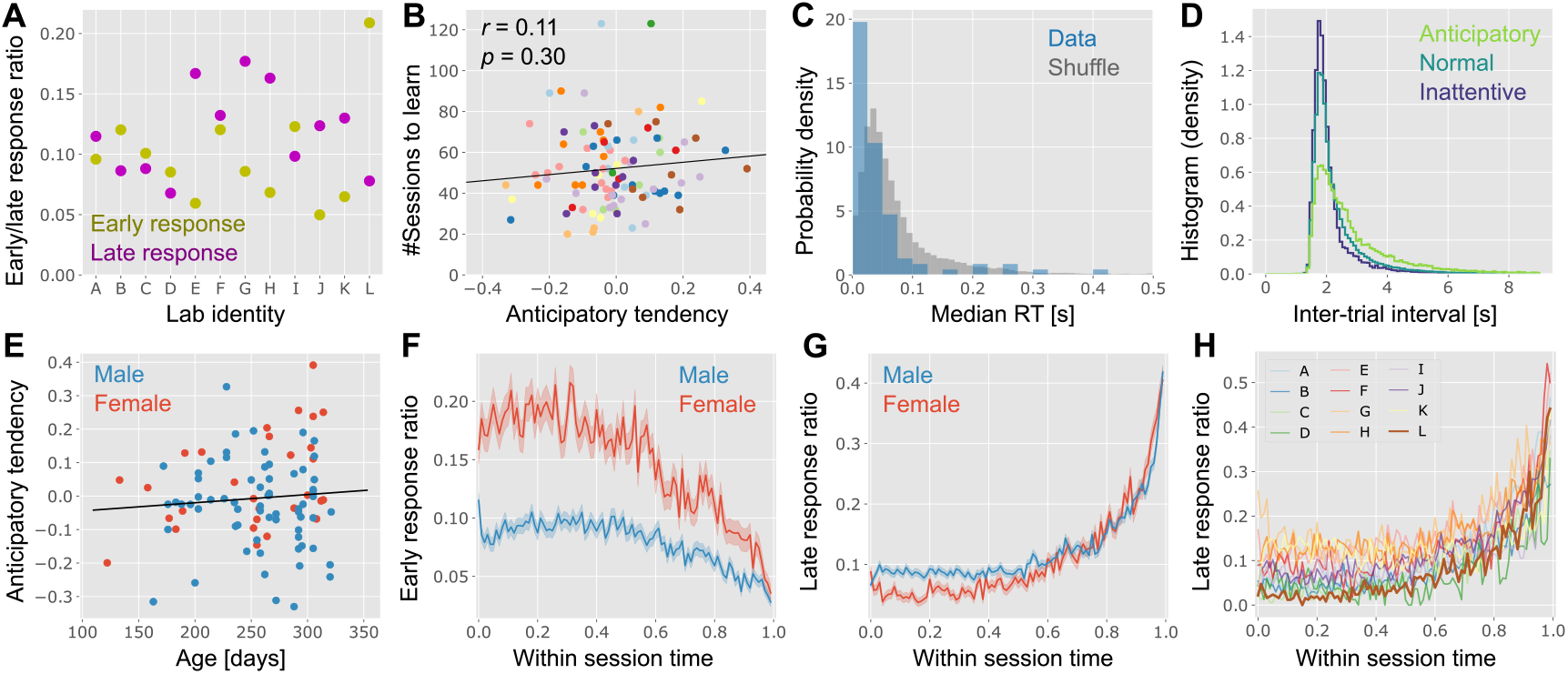
Reaction time variability. **(A)** Average early and late response ratios for each lab. As in Fig. 2E, lab L is an outlier. **(B)** The inter-animal variability in the task acquisition speed, measured by the total number of sessions an animal performed before starting an electrophysiology session (including habituation sessions), was not correlated with the anticipatory tendency. Points are color-coded by the lab identity as in Fig. 2E. **(C)** The standard deviation of the median RT across sessions for each mouse. The gray histogram is a shuffle estimated by randomly re-assigning sessions to animals. **(D)** The ITI distributions of anticipatory (ATI > 0.15), normal, inattentive mice (ATI < -0.15). As in Supp. Fig. S1H, we included the quiescent period into ITI. **(E)** Anticipatory tendency as a function of the age of mice. We did not see any significant trends; see Zang et al., (2025)^46^ for task performance by older (> 12 months) animals. **(F**,**G)** Sex difference in the within session behavioral dynamics. Early (F) and late (G) response ratio are plotted as a function of the relative position of a trial in a session. This is the same as Fig. 1G, but plotted male and female mice differently. **(H)** Lab identity dependence of the late response ratio as a function of the relative position of a trial in a session. Lab L (brown line) was plotted in a thick line for clarity.

**Supplementary Figure 3:**
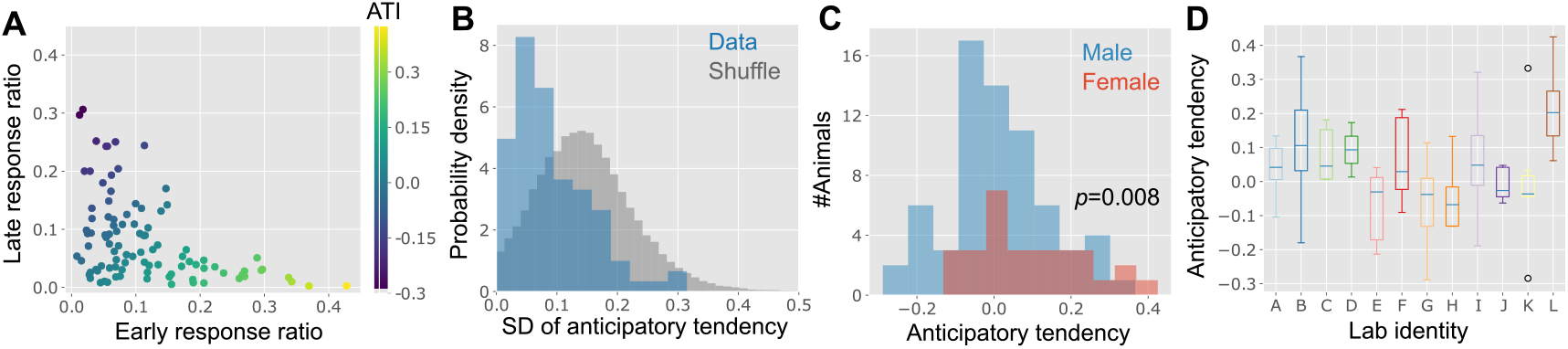
Behavioral variability statistics under a different cutoff. Instead of excluding last 40 trials as in the rest, here we excluded last 200 trials from the analysis to analyze the robustness of our main results. **(A)** Mouse-to-mouse variability in early/late response ratio. **(B)** The standard deviation (SD) of the anticipatory tendency over sessions calculated for each mouse, compared to random shuffle, as in Fig. 2B **(C)** Sex dependence of the anticipatory tendency (*t*(49.8) = 2.79, *p* = 0.008; Welch’s t-test on population means), **(D)** Relationship between the anticipatory tendency and lab identity. Note that panels A-D correspond to Figs. 2A, 2B, 2F, and 2E, respectively.

**Supplementary Figure 4:**
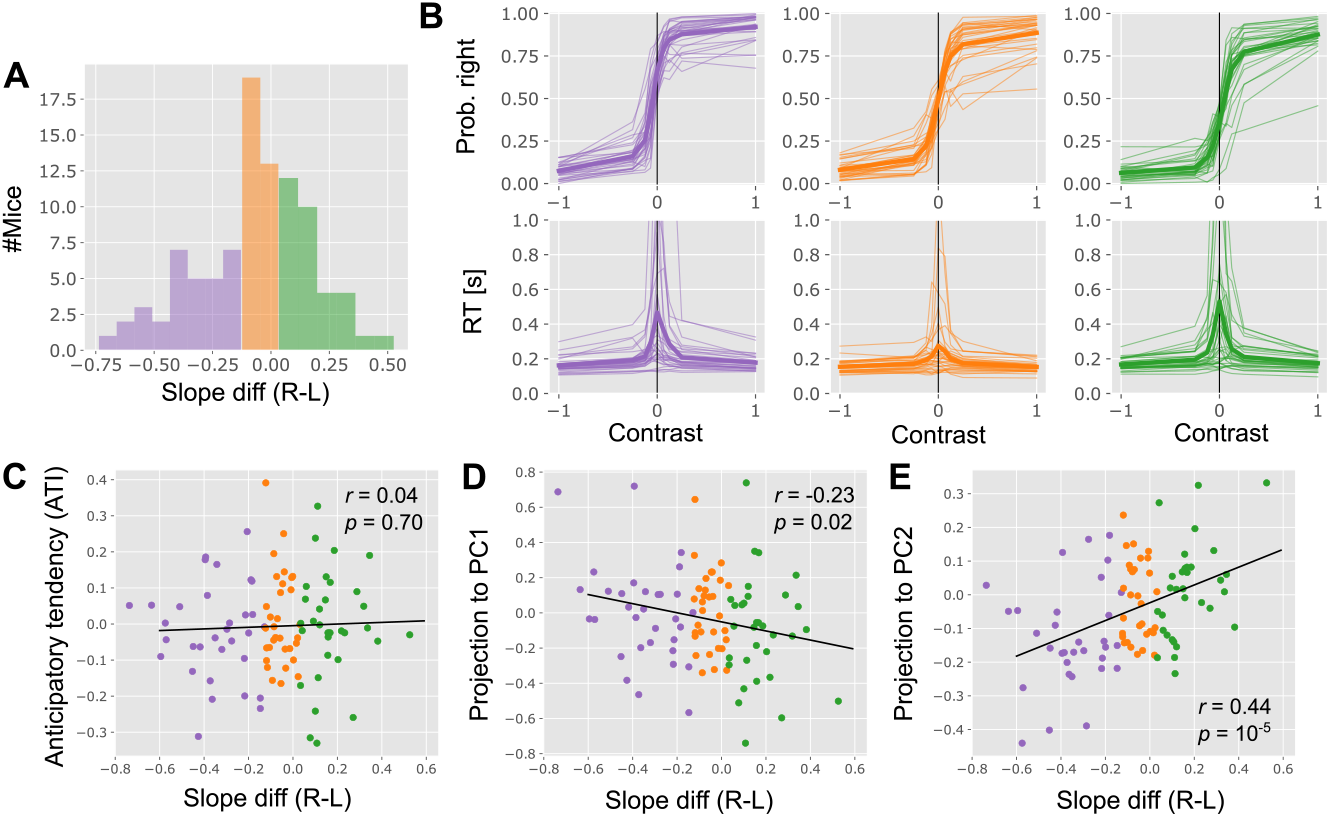
Analysis of individual variability based on psychometric curve slopes. **(A)** Histogram of left–right slope differences. Following Liebana et al.^6^, the left (right) slope was defined as the absolute difference between the probability of a rightward choice (defined by the final movement) under high-contrast (100% and 25%) left (right) stimulus conditions and the zero-stimulus condition. Slope difference was then defined as [right slope] - [left slope]. Data points were color-coded according to the 33rd and 67th percentiles. **(B)** Psychometric (top) and chronometric (bottom) curves for the three mouse groups defined in panel A. Thin lines represent individual mice, and the thick line indicates the group average. RT corresponds to the median of all valid RTs for each category. **(C–E)** Correlations between slope difference and anticipatory tendency (C), and projections onto PC1 (D) and PC2 (E) of the animal embedding vector. Slope difference showed a strong correlation with the PC2 projection and a weaker negative correlation with the PC1 projection, consistent with both dimensions being related to choice bias (Fig. 3I).

**Supplementary Figure 5:**
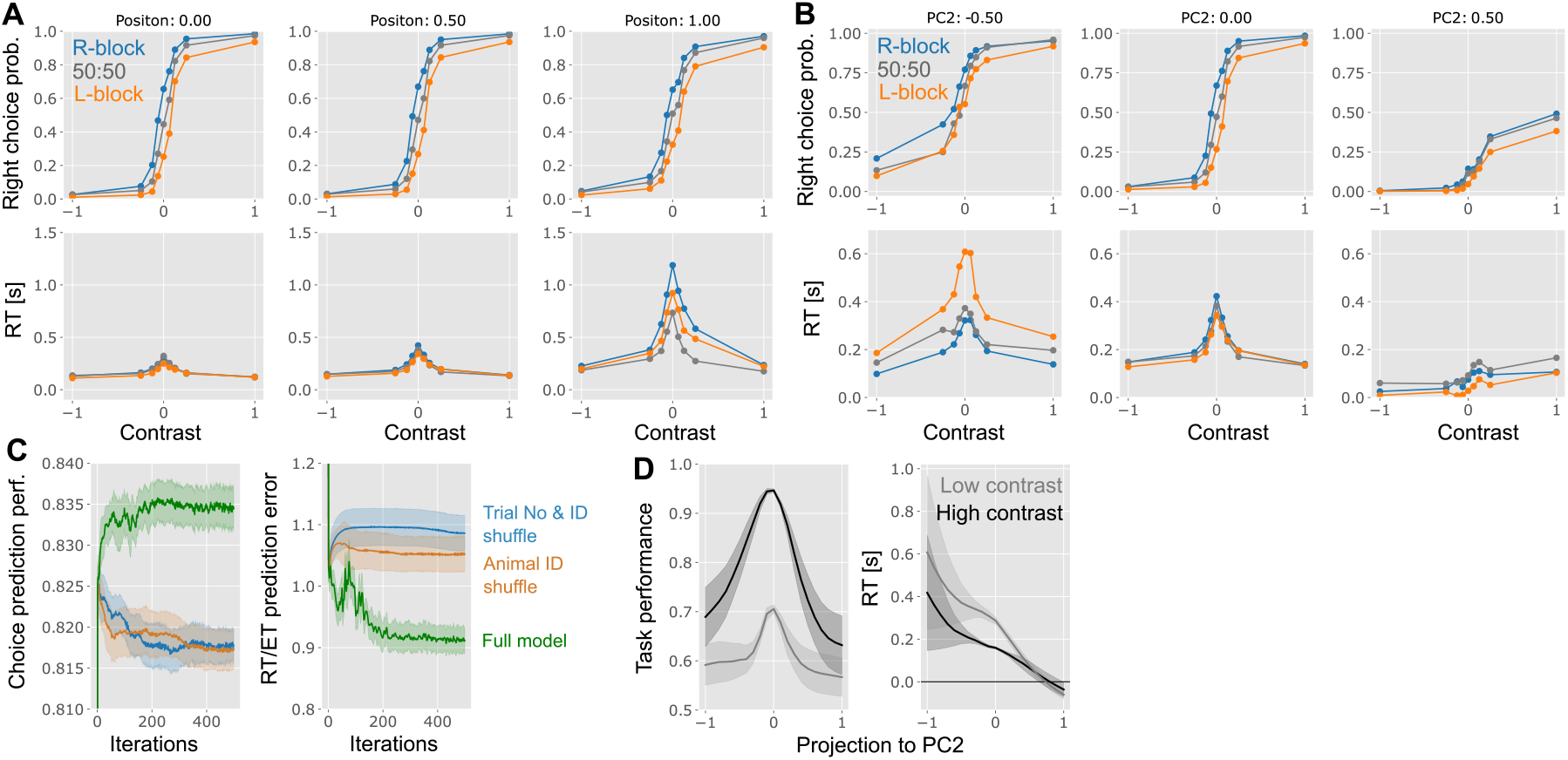
Psychometric and chronometric curves generated from animal-to-vec analysis. **(A)** Effect of within-task position on psychometric and chronometric curves. Positions 0, 0.5, and 1.0 correspond to the beginning, middle, and end of a session. **(B)** Shift in expected behavior along the second principal vector of the animal embedding. **(C)** Comparison of the model performance with shuffled models. We generated animal ID shuffling, depicted by the orange lines, by randomly shuffling the one-hot animal ID encoding across sessions. Similarly, in the blue curve, we shuffled both trial number encoding and animal IDs. **(D)** Average task performance and RT along the PC2 of the animal embedding.

**Supplementary Figure 6:**
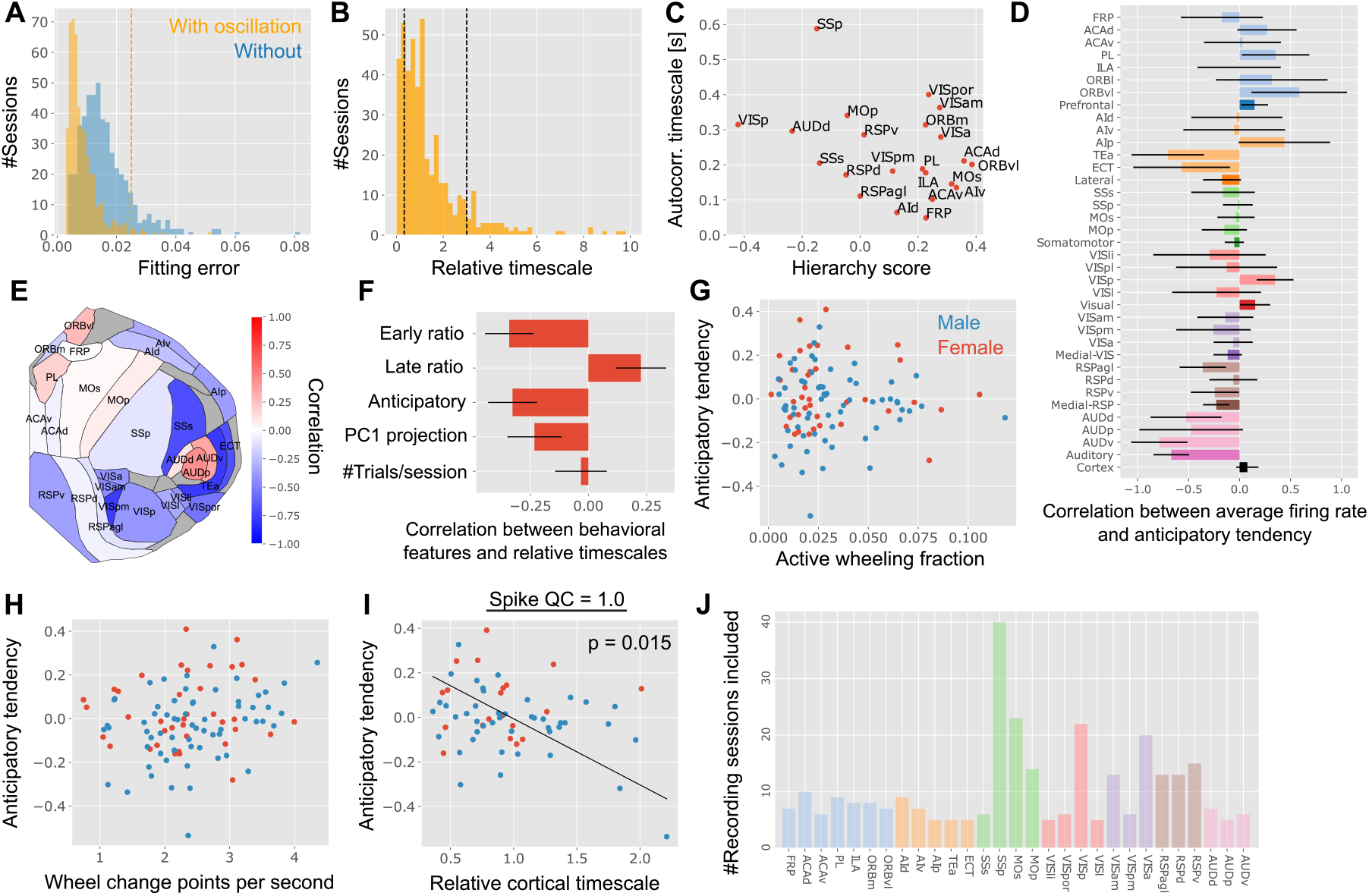
Supplementary analysis of passive period autocorrelation timescale. **(A)** The fitting error of the autocorrelation curve with or without oscillation components. The orange vertical line represents the cutoff for inclusion in the subsequent analysis. **(B)** The distribution of the estimated relative timescale. We excluded both overly small and large estimates from the analysis, as the inter-animal variance in autocorrelation is expectedly small. In panels A and B, if a session includes recording from multiple regions, they are counted independently. **(C)** Correlation between average firing rate of regions and divisions, and session-wise anticipatory tendency. **(D)** Estimated average cortical timescale and its relationship with the cortical hierarchy score^35^. **(E)** The same as Fig. 6D, but represented in a flattened cortical map. **(F)** Correlation between the relative cortical timescale of each animal and their behavioral statistics. We found a correlation with both early and late response ratios as they contribute equally to the anticipatory tendency. PC1 projection is the projection to the PC1 axis of the animal embedding. **(G, H)** Correlation between anticipatory tendency and the active fraction (*r* = − 0.03, *p* = 0.8; panel G) and the change points (*r* = 0.20, *p* = 0.04; panel H) of wheel trajectory during the passive period (see *Wheel statistics analysis* in Methods for the definition of wheel statistics). **(I)** The same as Fig. 6G, but only the neurons with the sorting quality criteria = 1.0 were used. For the analysis in the main text, we included all neurons regardless of the sorting quality, as we analyzed population-level autocorrelation. **(J)** Number of recording sessions included into the passive-period autocorelation analysis per region.

**Supplementary Figure 7:**
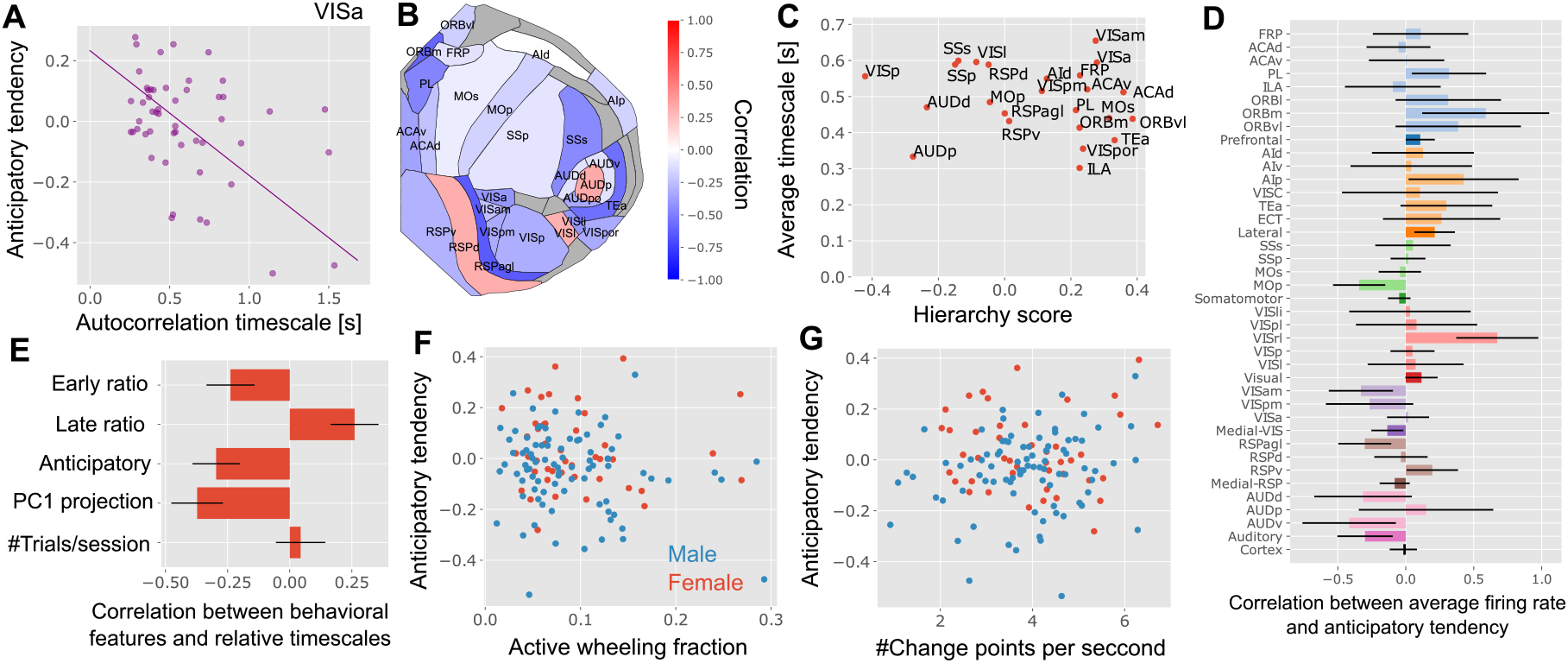
Supplementary analysis of neural autocorrelation and behaviors during ITI. **(A)** The correlation between the autocorrelation timescales at VISa and the session-wise anticipatory tendency (Pearson’s *r* = − 0.41, *p* = 0.003 without Bonferroni correction). Here, each point represents one session. **(B)** The same as Fig. 7D, but represented in a flattened cortical map. **(C)** Estimated average cortical timescale and its relationship with the hierarchy score^35^. **(D)** Correlation between average firing rate of regions and divisions, and session-wise anticipatory tendency. **(E)** Correlation between the relative cortical timescale of each animal and their behavioral statistics. **(F**,**G)** Correlation between anticipatory tendency and the active fraction (*r* = − 0.12, *p* = 0.16; panel F) and change points (*r* = 0.11, *p* = 0.2; panel G) of wheel trajectory during ITI.

**Supplementary Figure 8:**
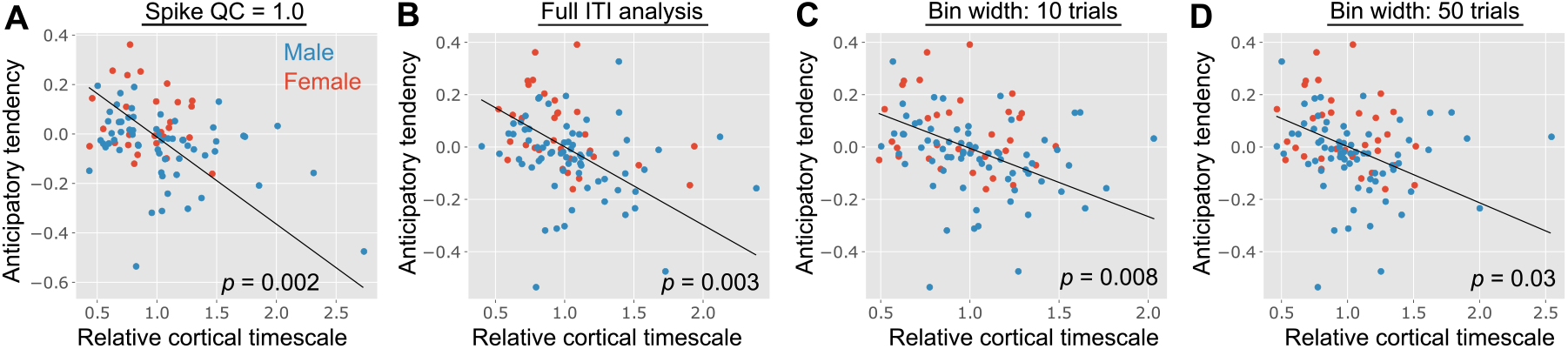
Correlation between animal-wise anticipatory tendency and relative cortical timescale under various model variants. **(A)** Only the neurons with spike-sorting quality = 1.0 are included, as opposed to QC = 0.0 we used in the main text. **(B)** The entire ITI period, defined as the duration between the beginning of ITI to 0.5 seconds before the stimulus onset, was used instead of the initial fraction of ITI. **(C, D)** The mean and variance of autocorrelation were estimated using 10 (C) and 50 (D) trial windows, instead of 30 trial windows in the main text.

**Supplementary Figure 9:**
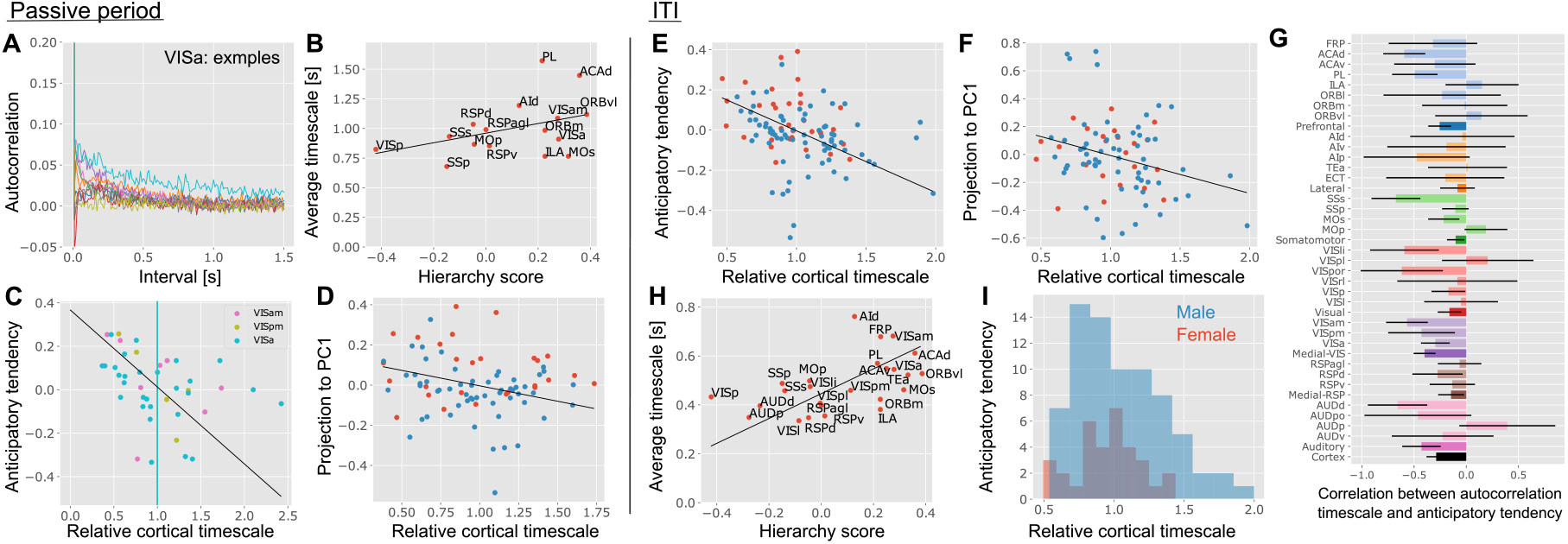
Neuron-wise fitting of autocorrelation. **(A)** Autocorrelation estimated from the passive period in a neuron-wise manner from a VISa recording. Each trace shows auto-correlation of one neuron. Please note that the scale of the y-axis is different from Figs. 6BC. **(B)** Average timescale of cortical regions as a function of hierarchy score (Pearson’s *r* = 0.41, *p* = 0.09, permutation test). **(C)** Correlation between session-wise anticipatory tendency and relative cortical timescale in medial visual areas (*r* = − 0.35, *p* = 0.018, *N*_session_ = 41, permutation test). Each point represents one session. **(D)** Animal-wise comparison between relative cortical timescale and the anticipatory tendency (*r* = − 0.15, *p* = 0.16, *N* = 84, permutation test). Each point represents one animal, color-coded by sex. **(E**,**F)** Correlation between the relative cortical timescale and the animal-wise anticipatory tendency (F; *r* = − 0.301, *p* = 0.002; permutation test) and the projection to PC1 of the animal embedding (G; *r* = − 0.27, *p* = 0.011; permutation-test). Each point represents one animal. **(G)** Correlation between average autocorrelation timescale of regions and divisions, and session-wise anticipatory tendency. **(H)** Correlation between the average timescale of each region and their hierarchy score (*r* = 0.51, *p* = 0.006; permutation test). **(I)** Relative cortical timescale of male and female was not statistically significant (*D* = 0.23, *p* = 0.16; KS-test). Panels A-D depict the results from the passive period, while panels E-I depict the results from the ITIs.

**Supplementary Figure 10:**
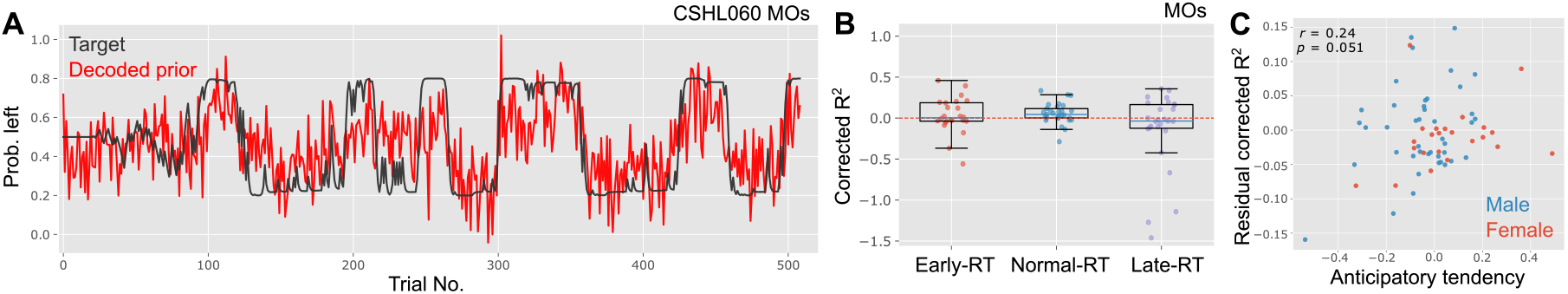
Prior-decoding from different RT types and animals with diverse ATI. **(A)** Example traces of the target Bayesian-optimal prior (black) and the prior decoded from secondary motor cortex (MOs) activity (red). We estimated both quantities following Findling et al.^28^ using the accompanying code (https://github.com/int-brain-lab/prior-localization/). For decoding, each neuron’s firing rate was computed from spiking activity in a 600–100 ms window before stimulus onset, and the prior was estimated via linear regression with an L1 regularizer^28^. **(B)** Corrected *R*^2^ between the target and decoded priors, computed separately for subsets of trials (early, normal, and late RT groups). Each point represents one session. We first decoded the prior using all trials within each session, then computed *R*^2^ for each trial group separately. To compensate for spurious correlations induced by slow autocorrelations, we subtracted the expected *R*^2^ under pseudo-block structure from the observed *R*^2 28,63^. The corrected *R*^2^ can be less than 1 due to this subtraction. For the early- and late-RT analyses, we included only sessions with more than 10 early-RT or late-RT trials, respectively. The mean corrected *R*^2^ under normal-RT was significantly larger than zero (*t*_stat_ = 2.47, *p* = 0.01; one-sample t-test). **(C)** Relationship between animal-wise anticipatory tendency and the residual corrected *R*^2^ of prior decoding (Pearson’s *r* = 0.24, *p* = 0.051). Each point represents one animal. Let 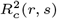 denote the corrected *R*^2^ for region *r* in subject *s*. We defined the mean residual corrected *R*^2^ for subject *s* as res- 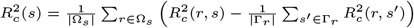, where Ω_*s*_ is the set of regions recorded in subject *s* and Γ_*r*_ is the set of subjects recorded in region *r*. For this analysis, we included only cortical regions that exhibited significant prior encoding in the previous work^28^. Animal-level anticipatory tendency was estimated as the average of session-wise anticipatory tendency over sessions included in the analysis. Both in panels B an C, only regions with equal to or greater than 10 neurons with QC=1.0 are included and the last 40 trials were excluded from corrected *R*^2^ estimation.

## SUPPLEMENTARY TEXT: BIAS IN AUTOCORRELATION ESTIMATION

Here, we prove that autocorrelation estimation from a short interval is biased and the proposed method mitigates this bias, assuming that ITI activity is sampled independently from an Gaussian Process. Let us consider estimation of the autocorrelation from a set of *M* time series, each containing observation at *t* = *t*_1_,.., *t*_*N*_. We generate a set of *M* time series *x*_*i*_(*t*_1_), …, *x*_*i*_(*t*_*N*_) by a Gaussian Process with kernel function *k*(·,·):

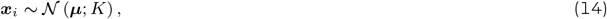

where ***x***_*i*_ ≡ [*x*(*t*_1_), …, *x*(*t*_*N*_)]. ***µ*** is an *N* -dimensional vector where all the elements are *µ*, and *K* is an *N × N* covariance matrix whose (*i, j*)-th element is *k*(*t*_*i*_, *t*_*j*_). Below, we restrict kernel function to stationary kernel (i.e., *k*(*t*_*i*_, *t*_*j*_) = *k*(*t*_*i*_ − *t*_*j*_)) for clarity, but the same argument holds for non-stationary kernels as well.

Suppose *t*_1_,.., *t*_*N*_ are sampled with the equal interval Δ*t* (i.e., *t*_*k*_ = *k*Δ*t*). Then, empirical estimation of auto-covariance for a single time series ***x*** is commonly defined by

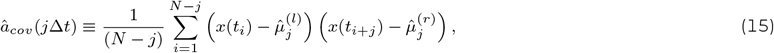

where the empirical left and right means are defined by

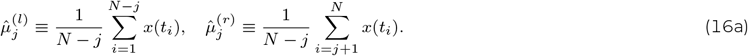

Below, we analyze autocovariance rather than autocorrelation for simplicity.

Let us first demonstrate that the above estimation of autocovariance is biased even at *M* → ∞ limit. For the ease of notation, we define 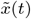 as the zero-meaned *x*(*t*):

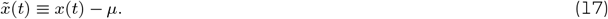

Taking *M* → ∞ limit and considering the expectation of â_*cov*_ (*j*Δ*t*) over the Gaussian process, we get

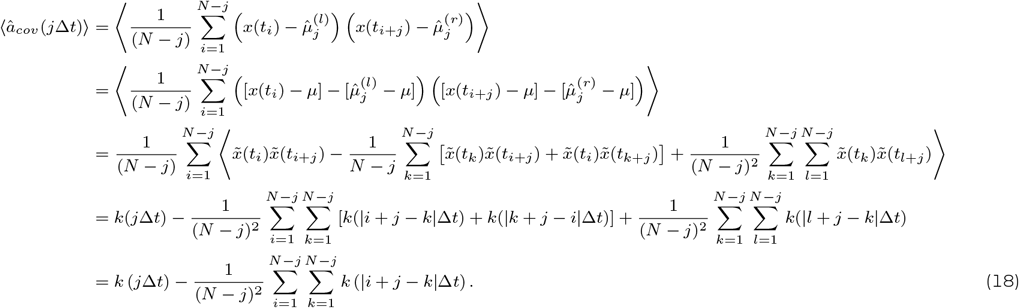

The equation above clearly shows that the empirical estimate of autocovariance â_*cov*_ is biased against the true auto-covariance *k*(*j*Δ*t*). This bias also results in the biased estimate of autocorrelation (trial-wise estimate of variance induces further bias). The bias term is 𝒪 (1) when *k* (|*i* + *j* − *k*|Δ*t*) is non-zero for most (*i, j, k*) combination. On the other hand, if *N* is large, such that the kernel *k* (|*i* + *j* − *k*|Δ*t*) decays to zero for most (*i, j, k*) combination, then the bias scales with 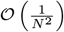.

A solution is to replace 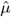 with the average activity over nearby trials. Specifically, we can define auto-covariance of the *µ*-th time series ***x***_*µ*_ = [*x*_*µ*_(*t*_1_), …, *x*_*µ*_(*t*_*N*_)] by

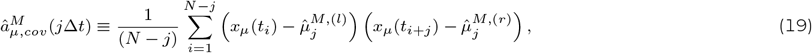

where the empirical left and right means are defined by

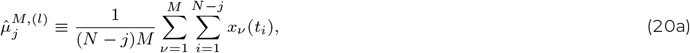

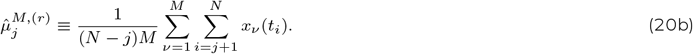

In this definition, from a parallel calculation with the one above, the expectation over GP is given by

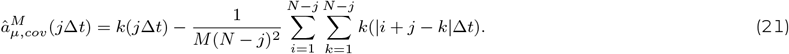

Thus, compared to Eq. 18, trial averaging reduces the bias by 1*/M* where *M* is the total number of trials used for the estimation of the mean.

This result indicates that it is best to use all the trials to minimize the bias. However, in the actual neural data, the i.i.d. assumption is violated. Thus, using all the trial may induce additional bias into the estimation of the mean activity. To navigate this balance, in the analysis, we used *M* = 30 for our analysis in the main result. Note that, by estimating the left and right means from different trials, we can further reduce the bias in principle, but it increases the variance in the estimate.

